# Genome sequencing of Syzygium cumini (Jamun) reveals adaptive evolution in secondary metabolism pathways associated with its medicinal properties

**DOI:** 10.1101/2023.07.12.548672

**Authors:** Abhisek Chakraborty, Shruti Mahajan, Manohar S. Bisht, Vineet K. Sharma

**Affiliations:** MetaBioSys Group, Department of Biological Sciences, Indian Institute of Science Education and Research Bhopal, Bhopal, India

**Author notes:** Corresponding Author email: Vineet K. Sharma. **E-mail addresses of authors:** Abhisek Chakraborty -, Shruti Mahajan -, Manohar S. Bisht -, Vineet K. Sharma.

## Abstract

Syzygium cumini, also known as jambolan or jamun, is an evergreen tree widely known for its medicinal properties, fruits, and ornamental value. To understand the genomic and evolutionary basis of its medicinal properties, we sequenced S. cumini genome, which is the largest genome sequenced for the first time from the world’s largest tree genus Syzygium using Oxford Nanopore and 10x Genomics sequencing technologies. The tetraploid and highly heterozygous draft genome of S. cumini had a total size of 709.9 Mbp with 61,195 coding genes. The phylogenetic position of S. cumini was established using a comprehensive genome-wide analysis including species from 18 Eudicot plant orders. The existence of neopolyploidy in S. cumini was evident from the higher number of coding genes and expanded gene families compared to the other two sequenced species from this genus. Comparative evolutionary analyses showed the adaptive evolution of genes involved in the phenylpropanoid-flavonoid (PF) biosynthesis pathway and other secondary metabolites biosynthesis such as terpenoid and alkaloid in S. cumini, along with genes involved in stress tolerance mechanisms, which was also supported by leaf transcriptome data generated in this study. The adaptive evolution of secondary metabolism pathways is associated with the wide range of pharmacological properties, specifically the anti-diabetic property, of this species conferred by the bioactive compounds that act as nutraceutical agents in modern medicine.

## INTRODUCTION

Syzygium cumini, also known as jamun, jambolan, or black plum, is a tropical tree belonging to the Myrtaceae plant family. It is native to the Indian subcontinent and South-East Asia, and is known for its wide range of medicinal properties and typical purple-black berries [1]. Syzygium, the clove genus, is the world’s largest tree genus with 1,193 recognized species. They occupy various habitats, medium to large-sized, typically sub-canopy trees, and thus affect the ecosystems of a wide range of organisms [2]. Some of the other species from Syzygium genus are - S. caryophyllatum, S. aromaticum, S. aqueum, S. grande, S. myrtifolium, etc., which are used as spices or fruits in pharmacology and horticulture industry [2, 3].

cumini is an evergreen tree with 30 meters of height, 3.6 meters of girth, and up to 15 meters of bole, and can live more than 100 years [4]. This Syzygium species is widely cultivated in tropical countries for its edible fruit (“Jamun”), which has significant economic importance [2]. S. cumini was introduced in several tropical and sub-tropical regions of the world for its commercial applications, such as Southern Africa, West Indies, California, and Israel [5]. The purple-black colored fruits of S. cumini are rich in anthocyanin, polyphenol, and tannin content and possess high nutrient values and medicinal properties [6]. Besides this, other parts of the tree, such as wood, leaf, flower, seed, and bark also have various economic and medicinal properties [5].

All the plant parts of S. cumini have therapeutic properties, which are used in various treatments since the Ayurvedic era [4]. The extracts from seed, bark, fruit, leaf, and flower of this species contain various phytochemicals such as glucoside jambolin, flavonoids including anthocyanin, terpenoids, and alkaloids (e.g., jambosine), which confer medicinal properties including anti-allergic, anti-oxidant, anti-diarrhoeal, anti-microbial, anti-inflammatory, anti-cancer, and others [1, 4, 5, 7]. Specifically, the fruit seed extracts of S. cumini have well-known anti-diabetic properties conferred by the glucoside jambolin present in the seeds [4, 5]. Besides, the leaves and bark extracts of S. cumini also have anti-diabetic potential due to the presence of phyenylpropanoids [1, 8]. Further, S. cumini fruits also possess anti-hyperlipidemic, hepatoprotective, anti-ulcer, anti-arthritic, anti fertility, and anti-pyretic activities [1]. Different parts of this tree are also rich in compounds containing isoquercetin, myrecetin, kaemferol, and others [9].

Syzygium, the tree genus with the highest number of species, is characterized by rapid speciation events, which resulted in a wide range of ecological and morphological diversity within the genus. A previous study has indicated that an ancient pan-Myrtales Whole Genome Duplication (WGD) event might have contributed to the early stages of diversification in the Myrtales plant order [2]. However, whole genome sequencing of only two Syzygium species has been performed until now [2, 10], and the whole genome and transcriptome assembly of S. cumini was not available.

Therefore, in this study, the genome sequencing of S. cumini was performed using 10x Genomics linked reads and Oxford Nanopore long reads to assemble its nuclear genome (710 Mbp) and chloroplast genome (158 Kbp). We inferred S. cumini genome to be tetraploid in this study, whereas previous studies have also shown the existence of intraspecific polyploidy (ranging from 2x to 6x) in S. cumini species [11, 12]. The phylogenetic position of S. cumini was resolved with respect to 17 other Eudicot orders, and comparative evolutionary analyses showed the key plant secondary metabolism pathways, such as the phenylpropanoid-flavonoid (PF) biosynthesis pathway, were adaptively evolved in this Syzygium species, which are responsible for the immense medicinal properties of this tree.

## METHODS

### Genome and transcriptome sequencing

The clean leaves of S. cumini were used for DNA-RNA extraction and species identification (Figure 1). DNA extraction was performed using Carlson lysis buffer (Supplementary Text). Quality check and quantification of the extracted DNA were carried out using Nanodrop 8000 spectrophotometer and Qubit 2.0 fluorometer, respectively. Species identification was performed using amplification and Sanger sequencing of the two marker genes - ITS2 (Internal Transcribed Spacer) and MatK (Maturase K), followed by BLASTN of the gene sequences against NCBI non-redundant (nt) database (Supplementary Text). The extracted DNA was used to prepare the 10x Genomics library on the Chromium controller instrument using Chromium Genome Library and Gel Bead Kit (10x Genomics), and sequenced on Illumina NovaSeq 6000 instrument. Further, the DNA was purified using Ampure XP magnetic beads (Beckman Coulter, USA), which was used to prepare the Nanopore library with SQK-LSK109 and SQK-LSK110 library preparation kit (Oxford Nanopore Technologies, UK) for sequencing on a MinION Mk1C sequencer.

**Figure 1.**
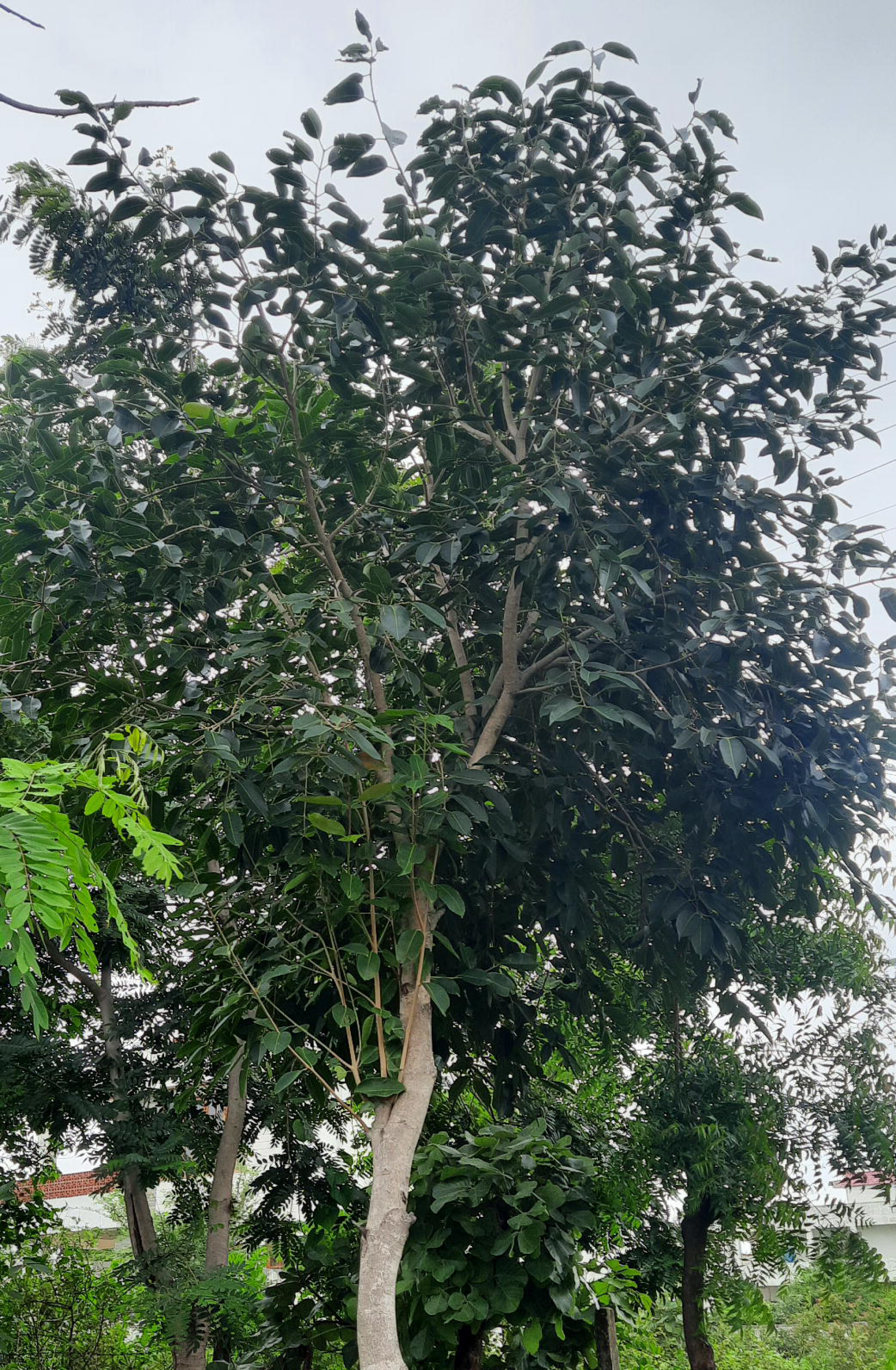
Syzygium cumini tree that was used for genome sequencing in this study.

The RNA was extracted following a similar method that was used for Syzygium longifolium species with a few modifications (Supplementary Text) [13]. Extracted RNA was washed and purified using a RNeasy mini kit (Qiagen, CA, USA). The RNA quality was diluted ten times and Qubit 2.0 fluorometer was used for quantification using a qubit ss RNA HS kit (Life Technologies, United States). Quality of the RNA was evaluated using High Sensitivity D1000 ScreenTape on Agilent 2200 TapeStation (Agilent, Santa Clara, CA). The RNA library was prepared using TruSeq Stranded Total RNA Library Preparation kit with the Ribo-Zero Plant workflow (Illumina Inc., CA, USA). The transcriptome library was sequenced on Illumina NovaSeq 6000 instrument to generate 150 bp paired-end reads.

### Genome assembly

Genomic characteristics such as genome size, genomic ploidy, and heterozygosity content were predicted using the 10x Genomics short reads before genome assembly. For this, barcode sequences were filtered using proc10xG (https://github.com/ucdavis-bioinformatics/proc10xG). The barcode filtered reads were further processed using Trimmomatic v0.39 with the filtering parameters - ‘‘TRAILING:20 LEADING:20 MINLEN:60 SLIDINGWINDOW:4:20” [14]. The filtered short reads were used to estimate the ploidy level of S. cumini genome using Smudgeplot v0.2.2 [15]. Further, these pre-processed reads were used to construct the k-mer count-based histogram with Jellyfish v2.2.10 [16], which was used to predict the genome size and heterozygosity content with GenomeScope v2 [15].

Oxford Nanopore long reads were processed to remove adapter sequences using Porechop v0.2.4, which were further error-corrected and de novo assembled using Canu v2.2 [17]. The resultant genome assembly was polished thrice with Pilon v1.23 using the pre-processed 10x Genomics short reads to fix any assembly errors [18]. Scaffolding was performed with the quality-filtered RNA-Seq reads (filtered using Trimmomatic v0.39), barcode-processed 10x Genomics reads (processed using Longranger basic v2.2.0), and error-corrected Nanopore reads using AGOUTI v0.3, ARCS v1.1.2, and LINKS v1.8.6, respectively [19–21]. Gap-closing of this scaffolded assembly was carried out using the error-corrected Nanopore long reads using LR_Gapcloser (three iterations) [22]. Finally, the pre processed 10x Genomics reads were used to again polish the assembly using Pilon v1.23 [18], and scaffolds with lengths of ≥5 Kb were retained to construct the final genome assembly of S. cumini.

After constructing the final genome assembly, the genomic ploidy level was further verified using nQuire [23]. The pre-processed linked reads were mapped onto the assembled genome using BWA-MEM v0.7.17 [24], and using these alignments, base frequencies were modeled using a Gaussian Mixture Model in nQuire. Log-likelihood values were estimated for each fixed model using the denoised base frequency distribution. The fixed model with the lowest Δlog-likelihood value compared to the free model was considered as the predicted ploidy level.

To assess the genome assembly quality, the pre-processed 10x Genomics linked reads, error corrected Nanopore reads and quality-filtered RNA-Seq reads were mapped onto the assembled genome to calculate the read mapping percentage using BWA-MEM v0.7.17 [24], Minimap v2.17 [25], and HISAT v2.2.1 [26], respectively. BUSCO v5.4.4 was used to check the presence of single copy orthologous genes in the final genome assembly with embryophyta_odb10 dataset [27]. Further, LTR Assembly Index (LAI) score was also calculated to evaluate the genome assembly quality using GenomeTools v1.6.1 and LTR_retriever v2.9.0 [28, 29].

To identify sequence variation in the S. cumini genome assembly, the pre-processed linked reads were mapped using BWA-MEM followed by variant calling using BCFtools “mpileup” v1.9 with the parameters - depth ≥30, variant sites quality ≥30, and mapping quality ≥50 [24, 30].

The chloroplast genome of S. cumini species was assembled with the pre-processed 10x Genomics data using GetOrganelle v1.7.7.0 with embplant_pt as seed database [31]. The chloroplast genome was annotated using GeSeq in CHLOROBOX with chloroplast genomes of other Syzygium species (S. aromaticum, S. forrestii, S. jambos, and S. malaccense) available in NCBI RefSeq database as reference sequences [32].

### Genome annotation

The whole genome assembly of S. cumini was used for constructing a de novo repeat library with RepeatModeler v2.0.3 [33], which was used to soft-mask the S. cumini genome with RepeatMasker v4.1.2 (http://www.repeatmasker.org). The repeat-masked genome was used to identify the coding genes with MAKER v3.01.04 genome annotation pipeline using AUGUSTUS as the ab initio gene predictor [34, 35]. For evidence-based alignments, de novo transcriptome assembly of S. cumini constructed in this study using Trinity v2.14.0 [36], and protein sequences of the two sequenced species from Syzygium genus - S. aromaticum and S. grande [2, 10] and other species from Myrtales order (Eucalyptus grandis and Corymbia citriodora) available in Ensembl plants release 56 were used [37]. A high-confidence coding gene set was constructed with the filtering criteria of AED value <0.5 and coding gene length ≥150 nucleotides.

The completeness of this coding gene set was evaluated using BUSCO v5.4.4 with embryophyta_odb10 database [27]. Gene expression values (TPM) were also estimated by mapping the quality-filtered RNA-Seq data of S. cumini onto the coding gene set (nucleotides) using Kallisto v0.48.0 [38].

The genome assembly of S. cumini was used for prediction of non-coding RNAs. de novo prediction of rRNA and tRNA was performed using Barrnap v0.9 (https://github.com/tseemann/barrnap) and tRNAscan-SE v2.0.9 [39], respectively. miRNA sequences were identified using BLASTN (sequence identity 80% and e-value 10^-9^) against the miRBase database [40].

### Collinearity analysis

MCScanX was used to analyse the intra-species collinearity for S. cumini species using the BLASTP homology alignments of coding genes and GFF annotations [41]. Further, inter-species collinear blocks were identified between S. cumini and S. grande, S. grande and S. aromaticum, and S. aromaticum and S. cumini using previously available data [2, 10]. Gene duplication analysis was also performed for the three Syzygium species using MCScanX.

### Phylogenetic analysis

For constructing the species phylogenetic tree, one Eudicot species from each available plant order (except Myrtales) in Ensembl plants release 56 was considered [37]. For Myrtales order, both the species available in Ensembl plants release 56 – E. grandis and C. citriodora were considered, along with S. grande and S. aromaticum from previous studies [2, 10]. The selected species from other 17 Eudicot plant orders are – Arabidopsis thaliana (Brassicales), Actinidia chinensis (Ericales), Beta vulgaris (Caryophyllales), Citrus clementina (Sapindales), Cucumis sativus (Cucurbitales), Coffea canephora (Gentianales), Cynara cardunculus (Asterales), Daucus carota (Apiales), Gossypium raimondii (Malvales), Juglans regia (Fagales), Kalanchoe fedtschenkoi (Saxifragales), Rosa chinensis (Rosales), Populus trichocarpa (Malpighiales), Sesamum indicum (Lamiales), Solanum tuberosum (Solanales), Vigna radiata (Fabales), and Vitis vinifera (Vitales). Alongside, Zea mays was considered as an outgroup species.

Proteome files of these 23 species with the longest isoforms for each protein were used for orthogroups construction with OrthoFinder v2.5.4 [42]. The orthogroups were filtered to extract the fuzzy one-to-one orthogroups using KinFin v1.1 [43]. Only those orthogroups comprising sequences from all 23 species were considered, and each orthogroup was processed to include only one longest sequence per species. The resultant orthogroups were individually aligned using MAFFT v7.310 [44], and filtered and concatenated with BeforePhylo v0.9.0 (https://github.com/qiyunzhu/BeforePhylo). The species phylogenetic tree was constructed with this concatenated alignment using maximum likelihood-based RAxML v8.2.12 with 100 bootstrap values and “PROTGAMMAAUTO” model [45].

### Analysis of gene family evolution

Proteome files of 23 species with the longest isoforms for each protein were used to analyse the expansion/contraction of gene families with CAFÉ v5 [46]. Homology-based search result obtained from All-versus-All BLASTP was clustered and gene families were filtered as per the suggestions for performing CAFÉ analysis. An ultrametric species phylogenetic tree across the 23 species was constructed using the calibration point for S. cumini and B. vulgaris (118 years), as reported in the TimeTree database v5 [47]. The ultrametric species tree and the filtered gene families were used in the two-lambda (λ) model implemented in CAFÉ v5, where species from Myrtales order were indicated separate λ-value compared to the other species.

### Identification of evolutionary signatures in S. cumini genes

Comparative analysis was performed to identify evolutionary signatures in S. cumini genes across 13 Eudicot species including S. cumini. Four other species from Myrtales order itself (S. grande, S. aromaticum, E. grandis, and C. citriodora), and species from its closer plant orders were considered for the analysis. Species from other plant orders used in these analyses are – V. radiata (order Fabales), C. sativus (order Cucurbitales), J. regia (order Fagales), R. chinensis (order Rosales), P. trichocarpa (order Malpighiales), A. thaliana (order Brassicales), G. raimondii (order Malvales), and C. clementina (order Sapindales).

#### Unique amino acid substitution with functional impact

Protein sequences of the 13 species were used for orthogroups construction with OrthoFinder v2.5.4 [42]. Orthogroups comprising sequences from the 13 species were extracted, and each orthogroup was filtered to retain the longest sequence for each species. The resultant orthogroups were individually aligned with MAFFT v7.310, and from these multiple sequence alignments S. cumini genes were identified that showed different amino acids in positions where the other species had the same amino acid. In this analysis, gaps in the alignments and ten positions around the gaps were not considered. Further, impact of the unique substitutions on the protein function was predicted with SIFT using UniProt as a reference database [48].

#### Higher nucleotide divergence

The protein sequence alignments for the orthogroups obtained in the previous step were used for orthogroup-specific phylogenetic tree construction with RAxML v8.2.12 using 100 bootstrap values and “PROTGAMMAAUTO” model [45]. Using the gene phylogenetic trees, S. cumini genes showing greater branch length values compared to the genes from other species were identified using “adephylo” package in R, and were termed as the genes with higher nucleotide divergence [49].

#### Positive selection

The orthogroups constructed across 13 species (nucleotide sequences) were individually aligned with MAFFT v7.310 [44]. The resultant multiple sequence alignments and the species tree of 13 species (constructed with RAxML) were used to detect the positively selected genes in S. cumini with a branch-site model in “codeml” of PAML v4.10.6 [50]. Likelihood-ratio test (LRT) and chi-square analysis were performed, and genes qualifying against the null model with FDR-corrected p-values of < 0.05 were identified as the genes showing positive selection in S. cumini. Further, Bayes Empirical Bayes analysis was performed to detect the genes with codon sites under positive selection (with >95% probability) for the foreground lineage.

cumini genes with more than one of the evolutionary signatures – unique substitution with functional impact, positive selection, and higher nucleotide divergence were termed as the genes showing multiple signatures of adaptive evolution (MSA) [51, 52].

### Functional annotation

cumini coding gene set was annotated against publicly available databases - Swiss-Prot database using BLASTP (e-value 10^-5^), NCBI-nr database using BLASTP (e-value 10^-5^), and Pfam-A database using HMMER v3.1 (e-value 10^-5^) [53–56]. S. cumini coding gene set, including the expanded gene families and the genes with evolutionary signatures, were annotated using eggNOG-mapper v2.1.9 and KAAS v2.1 genome annotation servers [57, 58]. Over-representation analysis using WebGestalt web server was performed to assign Gene Ontology (GO) categories to the MSA genes of S. cumini [59].

### Gene structure analysis

Genes associated with phenylpropanoid-flavonoid (PF) biosynthesis pathway and terpenoid biosynthesis pathway were identified in S. cumini genome from the functional annotation of the coding genes. Gene families were identified from the CAFÉ analysis, and the longest gene for each gene family was extracted. The genes were mapped separately onto S. cumini genome constructed in this study and the previously available S. aromaticum [10] and S. grande [2] genomes using Exonerate v2.4.0 (https://github.com/nathanweeks/exonerate) to construct the exon-intron structures [60], and for a comparative analysis across the three Syzygium species.

## RESULTS

### Genome assembly

Species identification was performed using matK and ITS2 marker gene sequencing, which showed 99.65% and 99.7% sequence similarity (the best hits), respectively, with S. cumini gene sequences available in NCBI non-redundant nucleotide (nt) database. 120.7 Gb of 10x Genomics data and 14.4 Gb Oxford Nanopore data (read N50 = 10.9 Kb) were generated for genome assembly. Based on the predicted genome size of 730.3 Mbp (using GenomeScope), the genomic data corresponded to 165.3x and 19.7x sequencing coverage for 10x Genomics and Nanopore reads, respectively (Supplementary Table 1). Additionally, 15.1 Gb RNA-Seq data was also sequenced from the S. cumini leaf tissue.

cumini genome contained 3.25% heterozygosity (Supplementary Figure 1) and was inferred as a tetraploid genome since the distribution of base frequencies at the variable sites showed the smallest Δlog-likelihood value for the tetraploid fixed model (Supplementary Figure 2A). The heterozygous k-mer pair distribution showed that 87% of the k-mers represented the total coverage of k-mer pair 4n (Supplementary Figure 2B) [15].
cumini genome assembly had a size of 709.9 Mbp containing 7,702 scaffolds with an N50 value of 179.2 Kb, and the largest scaffold size of 1.6 Mb (Supplementary Table 2). S. cumini genome showed the presence of 98.3% complete BUSCOs (64.6% complete and single-copy, and 33.7% complete and duplicated) (Supplementary Table 3). The genome assembly also had an LAI (LTR Assembly Index) score of 11.69. Further, 97.83% of barcode-filtered 10x Genomics reads, 93.45% error-corrected Nanopore reads, and 95.25% quality-filtered RNA-Seq reads were mapped onto the genome assembly. A total of 6,184,849 base positions (0.87%) in the genome assembly had sequence variations.

The chloroplast genome assembly of S. cumini showed a circular genome of 158,509 bases with 86 protein-coding genes (Supplementary Figure 3). The assembled genome assembly and annotation statistics (Supplementary Table 4) were similar to the previously reported chloroplast genomes of S. cumini and other Syzygium species [61, 62].

### Genome annotation

A de novo repeat library consisting of 2,521 sequences for S. cumini genome was used to repeat mask 51.51% of the genome assembly. Among the repeat classes, 49.31% were interspersed repeats, including 8.09% Gypsy and 5.37% Copia elements (Supplementary Table 5). Using the repeat-masked genome assembly, coding genes were predicted in S. cumini genome using the MAKER pipeline [34].

A total of 204,525 transcripts were assembled with an N50 value of 2,313 bp (Supplementary Table 6), which were used as empirical evidence along with the protein sequences of species from Myrtales order during coding genes prediction. 74,657 coding genes were predicted, among which 62,971 were retained (84.35%) after AED-based filtering. Further, length-based filtering was performed to retain 61,195 coding genes in the final high-confidence gene set with an average CDS length of 1,106.2 bp. 32,888 of the coding genes (53.74%) showed relatively higher gene expression (TPM values > 1). Distribution of the coding genes in various KEGG pathways and COG categories are mentioned in Supplementary Tables 7-8.

This coding gene set showed the presence of 92.8% BUSCOs (78.7% complete and 14.1% fragmented) (Supplementary Table 3). 92.38% of the genes were annotated using any of the three publicly available databases – NCBI-nr, Pfam-A, and Swiss-Prot (Supplementary Table 9). 1,174 tRNAs decoding standard amino acids, 702 rRNAs, and 176 miRNAs were also identified in the assembled genome of S. cumini.

### Collinearity and orthologous gene clustering

Synteny analysis showed the presence of 16.76% intra-species collinearity in S. cumini genome. Further, 90.55% of the coding genes were originated by duplication events, whereas, gene duplication analysis using previously available data of Syzygium species showed a lesser percentage of duplicated genes in S. grande (75.94%) and S. aromaticum (85.06%) [2, 10]. Inter-species collinearity analysis showed a higher percentage of collinear genes between S. grande and S. aromaticum than S. cumini and S. grande, and S. cumini and S. aromaticum (Supplementary Table 10). Further, a higher percentage of S. cumini genes and a higher number of collinear blocks were present in the inter-species collinear blocks constructed between S. cumini and S. grande, compared to S. cumini and S. aromaticum (Supplementary Table 10). 17,882 S. cumini genes (29.22%) were present in the inter-species collinear blocks constructed with both S. aromaticum and S. grande, indicating their conserveness. The distribution of the 17,882 genes in KEGG pathways is mentioned in Supplementary Table 11.

Gene clustering among S. cumini and four other species from Myrtales order showed a large number of species-specific gene clusters in S. cumini (2,891 clusters) compared to other species (Figure 2). 3,980 gene clusters were common between S. cumini and S. grande, and 839 common gene clusters were identified between S. cumini and S. aromaticum. Genes included in the species-specific gene clusters of S. cumini (15,721 genes) were involved in various KEGG pathways mentioned in Supplementary Table 12.

**Figure 2.**
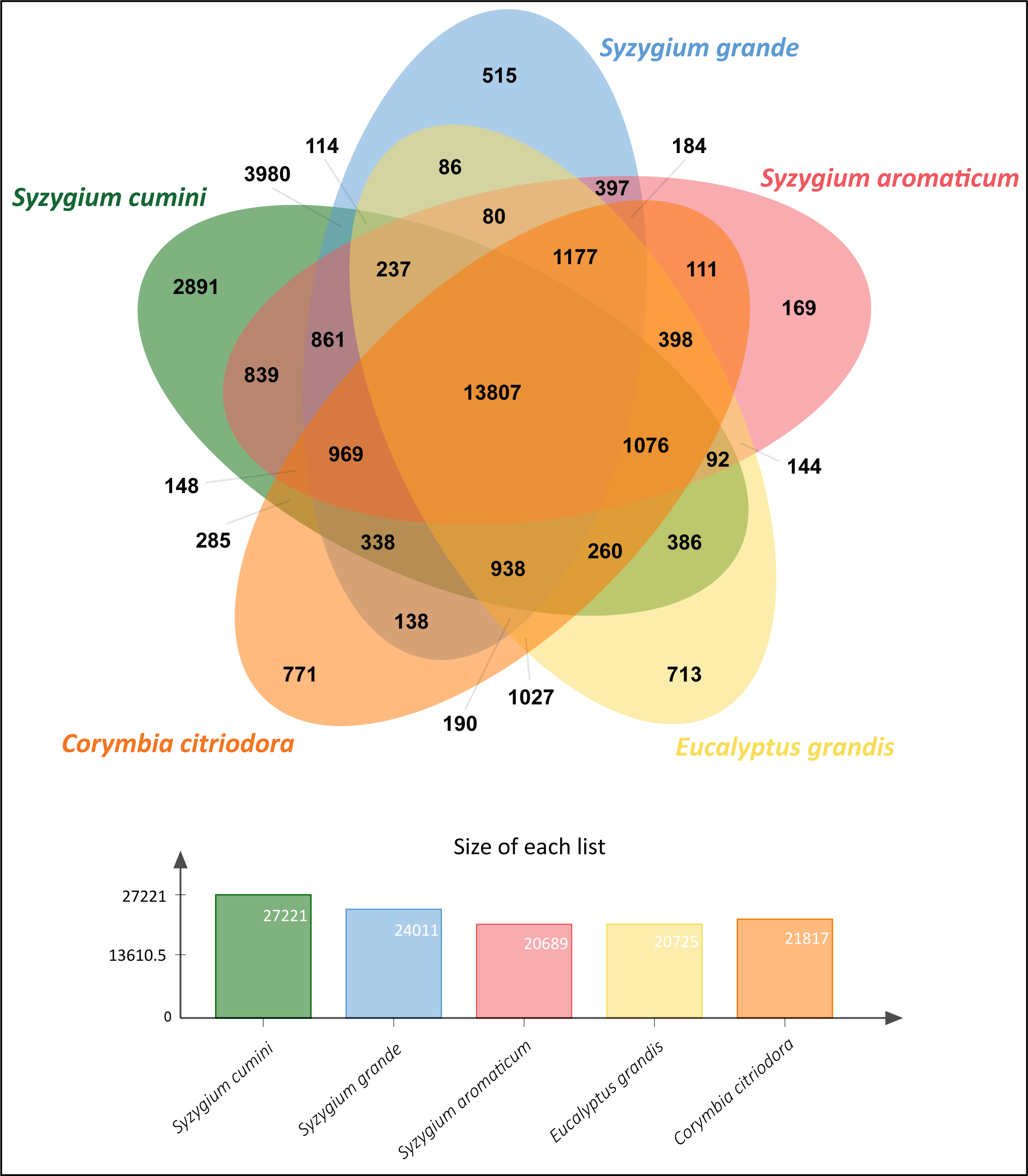
Orthologous gene clusters among S. cumini and other species from Myrtales plant order.

### Phylogenetic position of S. cumini

1,465 one-to-one fuzzy orthogroups were identified across 23 species spanning 18 Eudicot plant orders. Filtered and concatenated sequence alignments of the orthogroups containing 1,248,870 alignment positions were used to construct the species phylogenetic tree with Zea mays as the outgroup species.

In the phylogenetic tree, S. cumini was found in a position closer to S. grande (in the same clade) compared to S. aromaticum (Figure 3), which can further be explained by a higher number of collinear blocks and a higher number of shared gene clusters present between S. cumini and S. grande, compared to S. cumini and S. aromaticum (Figure 2, Supplementary Table 10). Among all the core Eudicot species in our phylogenetic tree, the species from the Saxifragales plant order (K. fedtschenkoi) diverged the earliest. The relative phylogenetic positions of the Eudicot orders were similar to the previous studies [63, 64].

**Figure 3.**
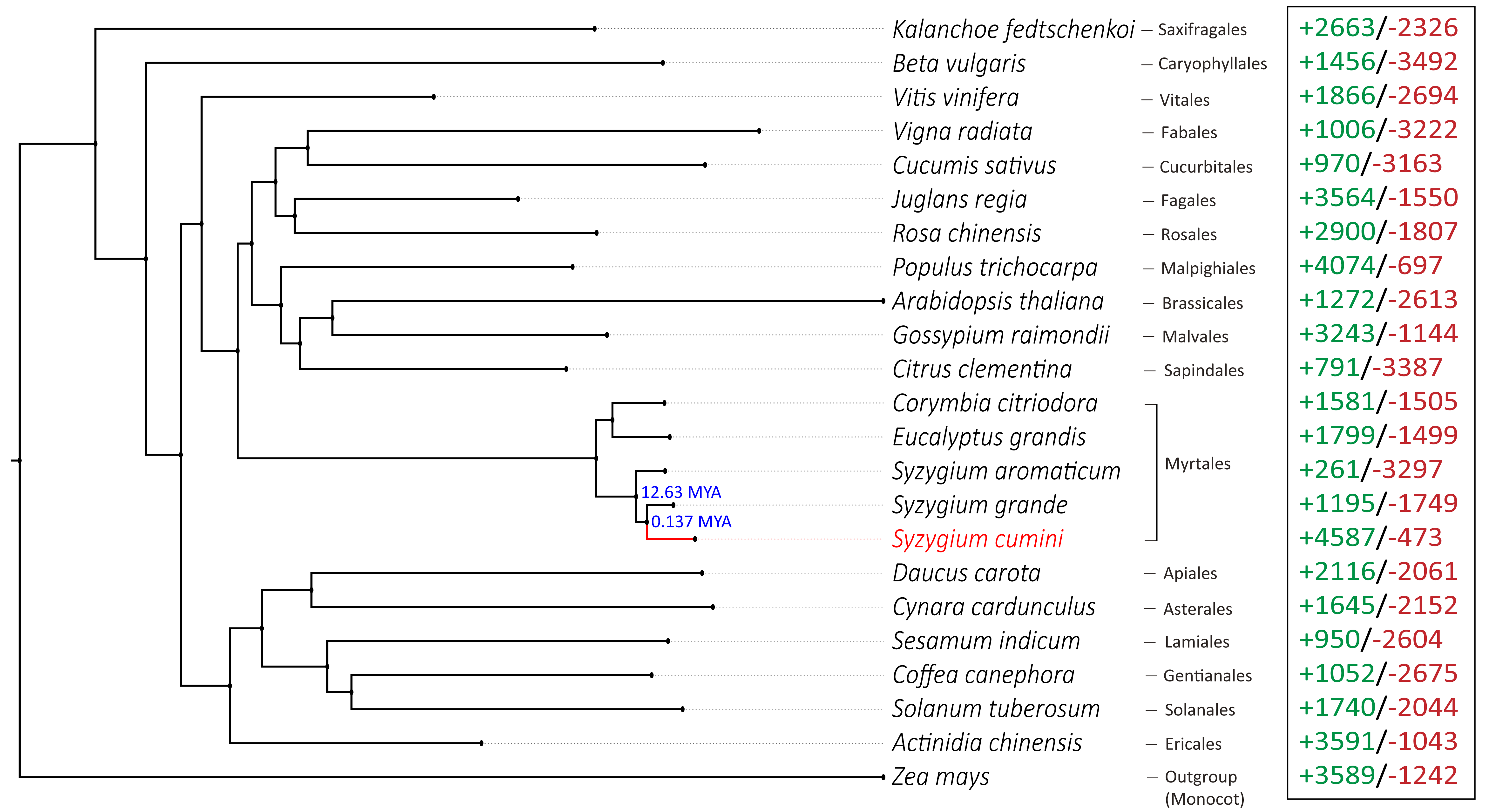
Phylogenetic position of S. cumini with respect to Eudicot species from Myrtales and 17 other plant orders. Zea mays was used as an outgroup species. Numbers mentioned in the nodes denote the divergence times of Syzygium species obtained from TimeTree v5 sdatabase [47]. Numbers in green and red represent the number of expanded and contracted gene families in each species, respectively.

### Gene family evolution

A total of 17,366 filtered gene families were identified across 23 species. Among these, 4,587 gene families were expanded, and 473 gene families were contracted in S. cumini species. The number of expanded gene families was much higher than that of S. grande and S. aromaticum (Figure 3). Among these expanded gene families, 41 families were highly expanded (>25 expanded genes) in S. cumini (Supplementary Table 13). The highly expanded gene families were involved in secondary metabolism-related pathways, such as Phenylpropanoid and flavonoid biosynthesis (Supplementary Table 14).

### Genes with evolutionary signatures

8,583 orthogroups across 13 species were constructed to identify the S. cumini genes with evolutionary signatures. 1,630 genes were positively selected (p-value < 0.05), 1,113 genes had unique amino acid substitution with functional impact, and 135 genes showed higher nucleotide divergence (Supplementary Tables 15-17). Among these genes, 430 genes had more than one signature of adaptive evolution (MSA genes). 333 of the MSA genes were also supported by gene expression (TPM > 1) data obtained in this study. GO categories of the S. cumini MSA genes are mentioned in Supplementary Table 18.

### Adaptive evolution of genes involved in secondary metabolism pathways

Plant secondary metabolites are derived from the primary metabolites and mainly function in the interaction of plants with their environment, abiotic and biotic stress tolerance, and are responsible for the medicinal properties of plants. The main classes of plant secondary metabolites are terpenoids, phenolic compounds, and alkaloids. The phenylpropanoid-flavonoid (PF) biosynthesis pathway is the key pathway for producing a wide range of phenolic compounds, such as flavonol, lignin, and anthocyanin [65]. Flavonoids and phenylpropanoids were the most abundant bioactive compounds in the fruit extracts of S. cumini, showing a wide range of pharmacological activities and can be used as preventive measures in many diseases, including type-2 diabetes [8].

#### PF biosynthesis pathway

Genes involved in the common phenylpropanoid pathway (conversion of phenylalanine to p-Coumaroyl CoA) showed gene family expansion, unique substitution with functional impact, and high gene expression (TPM > 1). p-Coumaroyl CoA, formed in the phenylpropanoid pathway, is also a precursor to flavonol, anthocyanin, and lignin biosynthesis [65, 66]. In the downstream phenylpropanoid pathway, genes showed gene family expansion/contraction, signatures of adaptive evolution, and high gene expression (TPM > 1).

#### Terpenoid biosynthesis pathway

Terpenoids are another important class of plant secondary metabolites. Fruits and flowers of S. cumini are rich in terpenoids [5, 8], and terpenoids present in the S. cumini leaves can be used to treat inflammatory diseases [67]. In support of this, seven key genes associated with terpenoid and other terpenoid-quinone biosynthesis showed adaptive evolution in S. cumini.

Using STRING database (v11.5) [68], protein-protein interaction was examined in the genes belonging to phenylpropanoid-flavonoid (PF) pathway and terpenoid biosynthesis pathway. Only the MSA genes (with TPM > 1) and the genes in highly expanded genes families of S. cumini were considered for the above analysis. Based on the protein-protein interaction evidence available on the STRING database, two clusters were formed by the terpenoid and phenolic compounds biosynthesis-related genes, and an association between these clusters was observed indicating a functional relationship between the two pathways.

#### Alkaloid and other secondary metabolites biosynthesis

The pharmacological activities of the alkaloids provide essential health benefits through the fruits and other plant parts of S. cumini [1]. Genes involved in isoquinoline and indole alkaloid biosynthesis pathways showed MSA (TPM > 1) and gene family expansion. Among other notable adaptively evolved genes, scopoletin, lignan, and glucoside biosynthesis-related genes were found.

### Adaptive evolution of genes associated with stress tolerance mechanisms

MSA genes of S. cumini were also involved in various biotic (such as pathogen resistance and defense against herbivores) and abiotic (such as ROS scavenging, heat, drought, and salinity, etc.) stress tolerance mechanisms. Among the key genes with MSA involved in biotic stress tolerance responses, GI downregulates salicylic acid accumulation and alters the phenylpropanoid pathway, thus reducing PR (Pathogenesis-Related) gene expression and negatively affecting biotic defense responses [69]. BSK provides resistance against bacterial and fungal pathogens by playing a role in pattern-triggered immunity (PTI) [70], NPR1 is a crucial regulator of salicylic acid signaling and triggers immune responses by inducing PR genes [71], MPK3 responds to biotic stress by upregulating jasmonic acid signaling and negatively regulating salicylic acid accumulation [72], PIK1 also acts in pathogen recognition and activation of defense responses [73].

Among the major genes with MSA involved in abiotic stress tolerance responses, ABF regulates the expression of abscisic acid-responsive genes to provide salinity, drought, and osmotic stress tolerance to plants [74], MPAO facilitates oxidative burst-mediated programmed cell death to aid plant defense responses [75], KUP K^+^ transporter family is involved in potassium deficiency and salt and drought stress response [76], Heat shock transcription factors (Hsf) regulates oxidative stress response by directly sensing the reactive oxygen species (ROS) [77]. Besides these, LOX confers abiotic (drought, salinity, etc.) and biotic stress tolerance [78], and CNGC has multifaceted functions in plants, such as pathogen resistance and abiotic (salt, drought, cold, etc.) stress tolerance [79].

## DISCUSSION

In this study, we performed whole genome sequencing of S. cumini species and constructed a draft genome assembly for the first time. It is only the third and till date the largest genome to be sequenced from the largest tree genus containing approximately 1,200 species. S. cumini was previously reported to show intraspecific polyploidy compared to S. aromaticum and S. grande [12]. Our analyses using two independent approaches to estimate the genomic ploidy also confirmed the tetraploidy in S. cumini genome. Further, the genome was found to be highly heterozygous (3.25%), and a combination of polyploidy and high heterozygosity increases the genomic complexity in this species. Polyploidy causes difficulty in haplotype resolving [80] and a higher percentage of allelic differences (1% or above) also poses a challenge in genome assembly [81]. Despite of this genomic complexity, we could successfully construct the whole genome assembly of S. cumini with the assembled genome size close to the predicted genome size.

We used multiple approaches such as BUSCO assessment, LAI score estimation, and read mapping percentage calculation to evaluate the genome assembly quality. 98.3% complete BUSCOs in the genome assembly suggest a near-complete genome assembly. LAI score of 11.69 indicates that the genome assembly constructed in this study can be considered as a “Reference” quality assembly [82]. LAI score of S. cumini genome constructed in this study was also similar to the other chromosome-scale plant genome assemblies such as Angelica sinensis [83]. A high percentage of mapped reads onto the genome assembly further attests to the assembly quality. Further, the usage of strict parameters, AED cut-off <0.5, and coding gene length >150 bp in the MAKER pipeline underscores the quality of the high-confidence coding genes. The presence of 92.8% BUSCOs in the coding gene set also suggests the near completeness of the genome annotation performed in this study. Further, the complete structures (exon-intron number and gene length) of the S. cumini genes involved in PF biosynthesis and terpenoid biosynthesis pathways could be identified, which was similar to the other two high-quality genome assemblies of Syzygium species, that also attests to the quality of S. cumini genome assembly.

We also noted a high percentage of complete and duplicated BUSCOs (D score) in the genome assembly and coding gene set of S. cumini (Supplementary Table 3), which is perhaps due to the additional neopolyploidy event following the Pan-Myrtales WGD event in S. cumini species (tetraploid) compared to the other Syzygium species that remained at the same ploidy level [2]. This event could also be the reason for an increased genome size of S. cumini compared to S. grande and S. aromaticum [2, 10]. The increase in genome size also appears to be due to an expansion in copy number (37% higher) of LTR-RT repeat elements in S. cumini genome compared to S. grande, and an overall 6.4% and 8.1% higher repeat content compared to S. grande and S. aromaticum genomes, respectively (Supplementary Table 5) [2, 10, 84]. The neopolyploidy event in S. cumini might also be the cause of a greater number of coding genes (61,195 genes), a greater fraction of genes (90.55%) originated from duplicated events, a higher number of gene clusters (27,221), and a higher number of species-specific gene clusters (2,891) observed in S. cumini species compared to S. grande and S. aromaticum (Figure 2). Duplicated genes may either undergo deletion or pseudogenization due to relaxed selection pressure [85] or acquire novel functions [86], which could also be the case in S. cumini species as observed in the adaptive evolution of secondary metabolism pathways.

Further, the consideration of species from Myrtales order (including two other Syzygium species) and species from its closer phylogenetic orders for comparative analysis to identify the genes with evolutionary signatures in S. cumini helped to reduce the false positives that could have resulted from the greater genetic distance of the selected species. The genes with evolutionary signatures identified from genomic analyses were also supported by transcriptomic analysis. It is important to note that MSA genes and the genes from highly expanded gene families were found to be majorly involved in secondary metabolites biosynthesis pathways such as phenylpropanoids, flavonoids, alkaloids, and terpenoids, which are responsible for the medicinal properties of this plant.

Phenylpropanoids play essential roles in plant development, response to abiotic and biotic stress signals, and biosynthesis of a broad spectrum of secondary metabolites [87]. Phenylpropanoid derived metabolites contribute to the biosynthesis of several other secondary metabolites, such as lignin and lignan, isoflavonoid, coumarin, stilbene, anthocyanin, isoquercetin, myrecetin, and kaemferol, which confer numerous pharmacological properties in S. cumini species [1, 5, 9]. One particular class of flavonoids - anthocyanin, is responsible for the purple-black color of the fruits of S. cumini and their health benefits [5]. Phenolic compounds (e.g., catechin, gallic acid, etc.) extracted from S. cumini seeds have immense potential as anti-diabetic and anti-oxidant agents, that have found commercial significance as nutraceutical ingredients in modern medicine and can be used as a substitution of allopathic remedies for chronic diseases such as type-2 diabetes [7, 88]. One of the main findings of this study was the identification of evolutionary signatures and gene family evolution of all the key S. cumini genes involved in the PF biosynthesis pathway. It is an important finding because the evolutionary signatures and evolution in gene families have been recognized as critical mechanisms shaping natural variation for species adaptation, which might also be the case in this species [52, 89]. Further, 741 genes were present in the expanded gene families of the PF biosynthesis pathway, among which 98.9% of the genes originated from different modes of duplication, which function in increasing the dosage of gene products and in accelerating the metabolic flux for rate-limiting steps in such biosynthetic pathways [90, 91]. Taken together, the adaptive evolution of PF biosynthesis pathway in S. cumini could be responsible for their numerous therapeutic properties, specifically the anti-diabetic property conferred by the seeds and leaves [1].

Notably, the comparative evolutionary analyses revealed seven key genes involved in the biosynthesis of terpenoids and other terpenoid-quinone compounds to show MSA and gene family expansion in S. cumini. Terpenoids are a structurally diverse class of secondary metabolites responsible for plant defence responses against herbivores and pathogens, and are abundant in S. cumini fruits contributing to the anti-oxidant and anti-inflammatory properties [92]. Other terpenoid-quinone compounds also function in plant stress tolerance responses [93] and show pharmacological activities [94]. Thus, the adaptive evolution of terpenoid biosynthesis pathway could explain the anti-inflammatory properties of S. cumini leaves and seeds conferred by the terpenoids [1, 67].

Among the other classes of secondary metabolites, alkaloids present in different plant parts of S. cumini are pharmaceutically diverse secondary metabolites with curative properties against many diseases [95]. Alkaloids, along with flavonoids and tannins also confer anti-arthritic property to the S. cumini seeds [1]. Glucosides are also critical secondary metabolites for plant defence responses and possess therapeutic properties [96]. S. cumini seed extracts contain alkaloid jambosine, and glucoside jambolin that prevents the conversion step of starch into sugar (anti-diabetic), which is the most significant therapeutic property of this species [9]. In this study, genes related to alkaloid and glucoside biosynthesis showed adaptive evolution in S. cumini that emphasizes the genomic basis for its pharmacological properties.

It is important to mention that the secondary metabolites are produced and regulated in response to various abiotic and biotic stresses, and aid in better survival of the plants and confer their medicinal properties [97]. Here, we also noted that various biotic and abiotic stress tolerance response genes displayed multiple signatures of adaptive evolution in S. cumini. Further, KEGG pathways related to phenylpropanoid, flavonoid, terpenoid, alkaloid biosynthesis, plant hormone signal transduction, and plant-pathogen interaction found in all the gene sets also showed the evolutionary signatures and gene family expansion (Supplementary Tables 14-17). Taken together, it is tempting to speculate that the adaptive evolution of major plant secondary metabolism pathways in S. cumini species confers unprecedented anti-diabetic, anti-oxidant, anti-inflammatory, and other pharmacological properties of this tree. Further, the whole genome sequence of S. cumini will facilitate future genomic, evolutionary, and ecological studies on the world’s largest tree genus.

## SUPPLEMENTAL INFORMATION

**Supplemental Information:** Supplementary Tables 1-18, Supplementary Figures 1-3.

## Acknowledgements

AC and SM thank Council of Scientific and Industrial Research (CSIR) for the research fellowship. MSB thanks Ministry of Education, Govt. of India for Prime Minister Research Fellowship (PMRF). The authors also thank the NGS facility at IISER Bhopal and the intramural research funds provided by IISER Bhopal.

## Authors’ contributions

VKS conceived and coordinated the project. SM prepared the samples for sequencing, performed Nanopore sequencing, and species identification. AC and VKS designed the computational framework of the study. AC performed all the computational analyses presented in the study and constructed all the figures. AC and MSB performed the functional annotation of gene sets. AC, VKS, and MSB interpreted the results. AC, VKS, and SM wrote the manuscript. All the authors have read and approved the final version of the manuscript.

## REFERENCES

1. Srivastava S, Chandra D. Pharmacological potentials of Syzygium cumini: a review. J Sci Food Agric. 2013;93:2084–93.

2. Low YW, Rajaraman S, Tomlin CM, Ahmad JA, Ardi WH, Armstrong K, et al. Genomic insights into rapid speciation within the world’s largest tree genus Syzygium. Nat Commun 2022 131. 2022;13:1–15.

3. Nair KN. The genus SyzygiumC: Syzygium cumini and other underutilized species.

4. Dagadkhair, AC. Jamun (Syzygium cumini) Skeels: A Traditional Therapeutic Tree and its Processed Food Products. Int J Pure Appl Biosci. 2017;5:1202–9.

5. Chaudhary B, Mukhopadhyay K. Syzygium cumini (L.) skeels: a potential source of nutraceuticals. Int J Pharm Biol Sci. 2012;2:46–53.

6. Ghosh P, Pradhan RC, Mishra S, Patel AS, Kar A. Physicochemical and nutritional characterization of jamun (Syzygium Cuminii). Curr Res Nutr Food Sci. 2017;5:25–35.

7. Kumar S, Sharma S, Kumar V, Sharma A, Kaur R, Saini R. Jamun (Syzygium cumini (L.) Skeels): The conventional underutilized multifunctional plant-an exotic gleam into its food and functional significance. Ind Crops Prod. 2023;191:115873.

8. Correia VT da V, Silva VDM, Mendonça H de OP, Ramos ALCC, Silva MR, Augusti R, et al. Efficiency of Different Solvents in the Extraction of Bioactive Compounds from Plinia cauliflora and Syzygium cumini Fruits as Evaluated by Paper Spray Mass Spectrometry. Mol 2023, Vol 28, Page 2359. 2023;28:2359.

9. Ayyanar M, Subash-Babu P. Syzygium cumini (L.) Skeels: A review of its phytochemical constituents and traditional uses. Asian Pac J Trop Biomed. 2012;2:240.

10. Ouadi S, Sierro N, Goepfert S, Bovet L, Glauser G, Vallat A, et al. The clove (Syzygium aromaticum) genome provides insights into the eugenol biosynthesis pathway. Commun Biol 2022 51. 2022;5:1–13.

11. Sharma N, Singh B, Wani MS, Gupta RC, Habeeb TH. Morphological, Cytological, and Chemotypic Variation of Essential Oils in Syzygium cumini (L.) Skeels. Anal Chem Lett. 2020;10:609–19.

12. Ohri D. How small and constrained is the genome size of angiosperm woody species. Silvae Genet. 2015;64:20–32.

13. Soewarto J, Hamelin C, Bocs S, Mournet P, Vignes H, Berger A, et al. Transcriptome data from three endemic Myrtaceae species from New Caledonia displaying contrasting responses to myrtle rust (Austropuccinia psidii). Data Br. 2019;22:794.

14. Bolger AM, Lohse M, Usadel B. Trimmomatic: A flexible trimmer for Illumina sequence data. Bioinformatics. 2014. https://doi.org/10.1093/bioinformatics/btu170.

15. Ranallo-Benavidez TR, Jaron KS, Schatz MC. GenomeScope 2.0 and Smudgeplot for reference free profiling of polyploid genomes. Nat Commun. 2020. https://doi.org/10.1038/s41467-020-14998-3.

16. Marçais G, Kingsford C. A fast, lock-free approach for efficient parallel counting of occurrences of k-mers. Bioinformatics. 2011. https://doi.org/10.1093/bioinformatics/btr011.

17. Koren S, Walenz BP, Berlin K, Miller JR, Bergman NH, Phillippy AM. Canu: Scalable and accurate long-read assembly via adaptive κ-mer weighting and repeat separation. Genome Res. 2017. https://doi.org/10.1101/gr.215087.116.

18. Walker BJ, Abeel T, Shea T, Priest M, Abouelliel A, Sakthikumar S, et al. Pilon: An integrated tool for comprehensive microbial variant detection and genome assembly improvement. PLoS One. 2014. https://doi.org/10.1371/journal.pone.0112963.

19. Zhang S V., Zhuo L, Hahn MW. AGOUTI: Improving genome assembly and annotation using transcriptome data. Gigascience. 2016. https://doi.org/10.1186/s13742-016-0136-3.

20. Yeo S, Coombe L, Warren RL, Chu J, Birol I. ARCS: Scaffolding genome drafts with linked reads. Bioinformatics. 2018. https://doi.org/10.1093/bioinformatics/btx675.

21. Warren RL, Yang C, Vandervalk BP, Behsaz B, Lagman A, Jones SJM, et al. LINKS: Scalable, alignment-free scaffolding of draft genomes with long reads. Gigascience. 2015. https://doi.org/10.1186/s13742-015-0076-3.

22. Xu GC, Xu TJ, Zhu R, Zhang Y, Li SQ, Wang HW, et al. LR-Gapcloser: A tiling path-based gap closer that uses long reads to complete genome assembly. Gigascience. 2018. https://doi.org/10.1093/gigascience/giy157.

23. Weib CL, Pais M, Cano LM, Kamoun S, Burbano HA. nQuire: A statistical framework for ploidy estimation using next generation sequencing. BMC Bioinformatics. 2018. https://doi.org/10.1186/s12859-018-2128-z.

24. Li H. Aligning sequence reads, clone sequences and assembly contigs with BWA-MEM. 2013;00:1–3.

25. Li H. Minimap2: Pairwise alignment for nucleotide sequences. Bioinformatics. 2018. https://doi.org/10.1093/bioinformatics/bty191.

26. Kim D, Langmead B, Salzberg SL. HISAT: A fast spliced aligner with low memory requirements. Nat Methods. 2015. https://doi.org/10.1038/nmeth.3317.

27. Simão FA, Waterhouse RM, Ioannidis P, Kriventseva E V., Zdobnov EM. BUSCO: Assessing genome assembly and annotation completeness with single-copy orthologs. Bioinformatics. 2015. https://doi.org/10.1093/bioinformatics/btv351.

28. Ou S, Jiang N. LTR_retriever: A highly accurate and sensitive program for identification of long terminal repeat retrotransposons. Plant Physiol. 2018. https://doi.org/10.1104/pp.17.01310.

29. Gremme G, Steinbiss S, Kurtz S. Genome tools: A comprehensive software library for efficient processing of structured genome annotations. IEEE/ACM Trans Comput Biol Bioinforma. 2013. https://doi.org/10.1109/TCBB.2013.68.

30. Danecek P, Bonfield JK, Liddle J, Marshall J, Ohan V, Pollard MO, et al. Twelve years of SAMtools and BCFtools. Gigascience. 2021;10:1–4.

31. Jin JJ, Yu W Bin, Yang JB, Song Y, Depamphilis CW, Yi TS, et al. GetOrganelle: A fast and versatile toolkit for accurate de novo assembly of organelle genomes. Genome Biol. 2020;21:1–31.

32. Greiner S, Lehwark P, Bock R. OrganellarGenomeDRAW (OGDRAW) version 1.3.1: Expanded toolkit for the graphical visualization of organellar genomes. Nucleic Acids Res. 2019;47:W59–64.

33. Flynn JM, Hubley R, Goubert C, Rosen J, Clark AG, Feschotte C, et al. RepeatModeler2 for automated genomic discovery of transposable element families. Proc Natl Acad Sci U S A. 2020. https://doi.org/10.1073/pnas.1921046117.

34. Campbell MS, Holt C, Moore B, Yandell M. Genome Annotation and Curation Using MAKER and MAKER-P. Curr Protoc Bioinforma. 2014. https://doi.org/10.1002/0471250953.bi0411s48.

35. Stanke M, Keller O, Gunduz I, Hayes A, Waack S, Morgenstern B. AUGUSTUS: A b initio prediction of alternative transcripts. Nucleic Acids Res. 2006. https://doi.org/10.1093/nar/gkl200.

36. Haas BJ, Papanicolaou A, Yassour M, Grabherr M, Blood PD, Bowden J, et al. De novo transcript sequence reconstruction from RNA-seq using the Trinity platform for reference generation and analysis. Nat Protoc. 2013. https://doi.org/10.1038/nprot.2013.084.

37. Bolser D, Staines DM, Pritchard E, Kersey P. Ensembl plants: Integrating tools for visualizing, mining, and analyzing plant genomics data. In: Methods in Molecular Biology. 2016.

38. Bray NL, Pimentel H, Melsted P, Pachter L. Near-optimal probabilistic RNA-seq quantification. Nat Biotechnol 2016 345. 2016;34:525–7.

39. Chan PP, Lin BY, Mak AJ, Lowe TM. TRNAscan-SE 2.0: Improved detection and functional classification of transfer RNA genes. Nucleic Acids Res. 2021. https://doi.org/10.1093/nar/gkab688.

40. Griffiths-Jones S, Saini HK, Van Dongen S, Enright AJ. miRBase: Tools for microRNA genomics. Nucleic Acids Res. 2008. https://doi.org/10.1093/nar/gkm952.

41. Wang Y, Tang H, Debarry JD, Tan X, Li J, Wang X, et al. MCScanX: A toolkit for detection and evolutionary analysis of gene synteny and collinearity. Nucleic Acids Res. 2012. https://doi.org/10.1093/nar/gkr1293.

42. Emms DM, Kelly S. OrthoFinder: Phylogenetic orthology inference for comparative genomics. Genome Biol. 2019. https://doi.org/10.1186/s13059-019-1832-y.

43. Laetsch DR, Blaxter ML. KinFin: Software for taxon-aware analysis of clustered protein sequences. G3 Genes, Genomes, Genet. 2017. https://doi.org/10.1534/g3.117.300233.

44. Katoh K, Standley DM. MAFFT multiple sequence alignment software version 7: Improvements in performance and usability. Mol Biol Evol. 2013. https://doi.org/10.1093/molbev/mst010.

45. Stamatakis A. RAxML version 8: A tool for phylogenetic analysis and post-analysis of large phylogenies. Bioinformatics. 2014. https://doi.org/10.1093/bioinformatics/btu033.

46. Mendes FK, Vanderpool D, Fulton B, Hahn MW. CAFE 5 models variation in evolutionary rates among gene families. Bioinformatics. 2020;36:5516–8.

47. Kumar S, Suleski M, Craig JM, Kasprowicz AE, Sanderford M, Li M, et al. TimeTree 5: An Expanded Resource for Species Divergence Times. Mol Biol Evol. 2022;39:1–6.

48. Ng PC, Henikoff S. SIFT: Predicting amino acid changes that affect protein function. Nucleic Acids Res. 2003. https://doi.org/10.1093/nar/gkg509.

49. Jombart T, Dray S. Adephylo: Exploratory Analyses for the Phylogenetic Comparative Method. Bioinformatics. 2010. https://doi.org/10.1093/bioinformatics/btq292.

50. Yang Z. PAML 4: Phylogenetic analysis by maximum likelihood. Mol Biol Evol. 2007. https://doi.org/10.1093/molbev/msm088.

51. Jaiswal SK, Mahajan S, Chakraborty A, Kumar S, Sharma VK. The genome sequence of Aloe vera reveals adaptive evolution of drought tolerance mechanisms. iScience. 2021. https://doi.org/10.1016/j.isci.2021.102079.

52. Chakraborty A, Mahajan S, Jaiswal SK, Sharma VK. Genome sequencing of turmeric provides evolutionary insights into its medicinal properties. Commun Biol 2021 41. 2021;4:1–12.

53. Bairoch A, Apweiler R. The SWISS-PROT protein sequence database and its supplement TrEMBL in 2000. Nucleic Acids Research. 2000.

54. Bateman A. The Pfam protein families database. Nucleic Acids Res. 2004. https://doi.org/10.1093/nar/gkh121.

55. Finn RD, Clements J, Eddy SR. HMMER web server: Interactive sequence similarity searching. Nucleic Acids Res. 2011. https://doi.org/10.1093/nar/gkr367.

56. Sharma AK, Gupta A, Kumar S, Dhakan DB, Sharma VK. Woods: A fast and accurate functional annotator and classifier of genomic and metagenomic sequences. Genomics. 2015;106:1–6.

57. Moriya Y, Itoh M, Okuda S, Yoshizawa AC, Kanehisa M. KAAS: An automatic genome annotation and pathway reconstruction server. Nucleic Acids Res. 2007. https://doi.org/10.1093/nar/gkm321.

58. Huerta-Cepas J, Forslund K, Coelho LP, Szklarczyk D, Jensen LJ, Von Mering C, et al. Fast genome wide functional annotation through orthology assignment by eggNOG-mapper. Mol Biol Evol. 2017. https://doi.org/10.1093/molbev/msx148.

59. Liao Y, Wang J, Jaehnig EJ, Shi Z, Zhang B. WebGestalt 2019: gene set analysis toolkit with revamped UIs and APIs. Nucleic Acids Res. 2019. https://doi.org/10.1093/nar/gkz401.

60. Chakraborty A, Bisht MS, Saxena R, Mahajan S, Pulikkan J, Sharma VK. Genome sequencing and de novo and reference-based genome assemblies of Bos indicus breeds. Genes Genomics 2023. 2023;1:1–10.

61. Asif H, Khan A, Iqbal A, Khan IA, Heinze B, Azim MK. The chloroplast genome sequence of Syzygium cumini (L.) and its relationship with other angiosperms. Tree Genet Genomes. 2013;9:867–77.

62. Tao L, Shi ZG, Long QY. Complete chloroplast genome sequence and phylogenetic analysis of Syzygium malaccense. http://www.tandfonline.com/action/authorSubmission?journalCode=tmdn20&page=instructions. 2020;5:3567–8.

63. Chakraborty A, Mahajan S, Bisht MS, Sharma VK. Genome sequencing and comparative analysis of Ficus benghalensis and Ficus religiosa species reveal evolutionary mechanisms of longevity. iScience. 2022;25.

64. Mahajan S, Chakraborty A, Sil T, Sharma VK. Genome sequencing and assembly of Tinospora cordifolia (Giloy) plant. bioRxiv. 2021;:2021.08.02.454741.

65. Taheri S, Teo CH, Heslop-Harrison JS, Schwarzacher T, Tan YS, Wee WY, et al. Genome Assembly and Analysis of the Flavonoid and Phenylpropanoid Biosynthetic Pathways in Fingerroot Ginger (Boesenbergia rotunda). Int J Mol Sci. 2022;23:7269.

66. Yadav V, Wang Z, Wei C, Amo A, Ahmed B, Yang X, et al. Phenylpropanoid Pathway Engineering: An Emerging Approach towards Plant Defense. Pathog 2020, Vol 9, Page 312. 2020;9:312.

67. Siani AC, Souza MC, Henriques MGMO, Ramos MFS. Anti-inflammatory activity of essential oils from Syzygium cumini and Psidium guajava. http://dx.doi.org/103109/138802092013768675. 2013;51:881–7.

68. Szklarczyk D, Gable AL, Nastou KC, Lyon D, Kirsch R, Pyysalo S, et al. The STRING database in 2021: customizable protein–protein networks, and functional characterization of user-uploaded gene/measurement sets. Nucleic Acids Res. 2021;49:D605–12.

69. Kundu P, Sahu R. GIGANTEA confers susceptibility to plants during spot blotch attack by regulating salicylic acid signalling pathway. Plant Physiol Biochem. 2021;167:349–57.

70. Majhi BB, Sreeramulu S, Sessa G. BRASSINOSTEROID-SIGNALING KINASE5 Associates with Immune Receptors and Is Required for Immune Responses. Plant Physiol. 2019;180:1166–84.

71. Chen J, Mohan R, Zhang Y, Li M, Chen H, Palmer IA, et al. NPR1 Promotes Its Own and Target Gene Expression in Plant Defense by Recruiting CDK8. Plant Physiol. 2019;181:289–304.

72. Jagodzik P, Tajdel-Zielinska M, Ciesla A, Marczak M, Ludwikow A. Mitogen-activated protein kinase cascades in plant hormone signaling. Frontiers in Plant Science. 2018.

73. Romeis T. Protein kinases in the plant defence response. Curr Opin Plant Biol. 2001;4:407–14.

74. Feng RJ, Ren MY, Lu LF, Peng M, Guan X, Zhou DB, et al. Involvement of abscisic acid-responsive element-binding factors in cassava (Manihot esculenta) dehydration stress response. Sci Reports 2019 91. 2019;9:1–12.

75. Yoda H, Hiroi Y, Sano H. Polyamine Oxidase Is One of the Key Elements for Oxidative Burst to Induce Programmed Cell Death in Tobacco Cultured Cells. Plant Physiol. 2006;142:193.

76. Yang T, Lu X, Wang Y, Xie Y, Ma J, Cheng X, et al. HAK/KUP/KT family potassium transporter genes are involved in potassium deficiency and stress responses in tea plants (Camellia sinensis L.): Expression and functional analysis. BMC Genomics. 2020;21:1–18.

77. Miller G, Mittler R. Could Heat Shock Transcription Factors Function as Hydrogen Peroxide Sensors in Plants? Ann Bot. 2006;98:279–88.

78. Viswanath KK, Varakumar P, Pamuru RR, Basha SJ, Mehta S, Rao AD. Plant Lipoxygenases and Their Role in Plant Physiology. J Plant Biol. 2020;63:83–95.

79. Guo J, Islam MA, Lin H, Ji C, Duan Y, Liu P, et al. Genome-wide identification of cyclic nucleotide gated ion channel gene family in wheat and functional analyses of TaCNGC14 and TaCNGC16. Front Plant Sci. 2018;9:18.

80. Kyriakidou M, Tai HH, Anglin NL, Ellis D, Strömvik M V. Current strategies of polyploid plant genome sequence assembly. Front Plant Sci. 2018;871:1660.

81. Asalone KC, Ryan KM, Yamadi M, Cohen AL, Farmer WG, George DJ, et al. Regional sequence expansion or collapse in heterozygous genome assemblies. PLoS Comput Biol. 2020. https://doi.org/10.1371/journal.pcbi.1008104.

82. Ou S, Chen J, Jiang N. Assessing genome assembly quality using the LTR Assembly Index (LAI). Nucleic Acids Res. 2018. https://doi.org/10.1093/nar/gky730.

83. Han X, Li C, Sun S, Ji J, Nie B, Maker G, et al. The chromosome-level genome of female ginseng (Angelica sinensis) provides insights into molecular mechanisms and evolution of coumarin biosynthesis. Plant J. 2022;112:1224–37.

84. Zhu S, Zhang X, Ren C, Xu X, Comes HP, Jiang W, et al. Chromosome-level reference genome of Tetrastigma hemsleyanum (Vitaceae) provides insights into genomic evolution and the biosynthesis of phenylpropanoids and flavonoids. Plant J. 2023. https://doi.org/10.1111/TPJ.16169.

85. Wang P, Moore BM, Panchy NL, Meng F, Lehti-Shiu MD, Shiu SH. Factors Influencing Gene Family Size Variation Among Related Species in a Plant Family, Solanaceae. Genome Biol Evol. 2018;10:2596.

86. Panchy N, Lehti-Shiu M, Shiu SH. Evolution of Gene Duplication in Plants. Plant Physiol. 2016;171:2294–316.

87. Vogt T. Phenylpropanoid Biosynthesis. Mol Plant. 2010;3:2–20.

88. Mahindrakar K V., Rathod VK. Antidiabetic potential evaluation of aqueous extract of waste Syzygium cumini seed kernel’s by in vitro α-amylase and α-glucosidase inhibition. Prep Biochem Biotechnol. 2021;51:589–98.

89. Guo YL. Gene family evolution in green plants with emphasis on the origination and evolution of Arabidopsis thaliana genes. Plant J. 2013;73:941–51.

90. Conant GC, Wolfe KH. Increased glycolytic flux as an outcome of whole-genome duplication in yeast. Mol Syst Biol. 2007;3:129.

91. Bekaert M, Edger PP, Chris Pires J, Conant GC. Two-Phase Resolution of Polyploidy in the Arabidopsis Metabolic Network Gives Rise to Relative and Absolute Dosage Constraints. Plant Cell. 2011;23:1719–28.

92. Cheng AX, Lou YG, Mao YB, Lu S, Wang LJ, Chen XY. Plant terpenoids: Biosynthesis and ecological functions. J Integr Plant Biol. 2007;49:179–86.

93. Liu M, Lu S. Plastoquinone and ubiquinone in plants: Biosynthesis, physiological function and metabolic engineering. Front Plant Sci. 2016;7:1–18.

94. Gordaliza M. Synthetic Strategies to Terpene Quinones/Hydroquinones. Mar Drugs. 2012;10:358.

95. Ziegler J, Facchini PJ. Alkaloid Biosynthesis: Metabolism and Trafficking. https://doi.org/101146/annurev.arplant59032607092730. 2008;59:735–69.

96. Bennett RN, Wallsgrove RM. Secondary metabolites in plant defence mechanisms. New Phytol. 1994;127:617–33.

97. Isah T. Stress and defense responses in plant secondary metabolites production. Biological research. 2019.

