## Supplemental Information for "Genome sequencing of Syzygium cumini (Jamun) reveals adaptive evolution in secondary metabolism pathways associated with its medicinal properties"

**Affiliation:**

*Corresponding Author email:

**E-mail addresses of authors:**

### SUPPLEMENTARY TABLES

**Supplementary Table 1. Summary of the raw genomic and transcriptomic data generated in this study for *S. cumini* species**

| Sequencing data | Total data (bases) | No. of reads | Average read length | Sequencing coverage* |
|---|---|---|---|---|
| Oxford Nanopore | 14,406,638,579 | 3,035,671 | 4,746 bases | 19.7X |
| 10x Genomics | 120,702,164,172 | 759,133,108 | 159 bases | 165.3X |
| RNA-Seq data | 15,098,162,682 | 94,956,998 | 159 bases | - |

*Sequencing coverage was calculated based on the estimated genome size of 730.3 Mbp

**Supplementary Table 2. Genome assembly statistics of *S. cumini* species (≥ 5 Kb)**

| Parameters | Values |
|---|---|
| No. of contigs | 7,702 |
| No. of contigs (≥ 10 Kb) | 7,558 |
| No. of contigs (≥ 25 Kb) | 6,160 |
| No. of contigs (≥ 50 Kb) | 3,642 |
| Total length | 709,916,230 |
| Total length (≥ 10 Kb) | 708,761,311 |
| Total length (≥ 25 Kb) | 683,107,220 |
| Total length (≥ 50 Kb) | 591,013,944 |
| Largest contig (bases) | 1,571,886 |
| N50 | 179,217 |
| L50 | 973 |
| GC% | 40.44 |
| N's per 100 kbp | 21.94 |

**Supplementary Table 3. BUSCO statistics of genome assembly and coding gene set**

| Parameters | Genome assembly | Coding gene set |
|---|---|---|
| Complete and single-copy (S) | 1,042 (64.6%) | 833 (51.6%) |
| Complete and duplicated (D) | 544 (33.7%) | 437 (27.1%) |
| Fragmented (F) | 9 (0.6%) | 228 (14.1%) |
| Missing (M) | 19 (1.1%) | 116 (7.2%) |
| Total BUSCO groups searched | 1,614 | 1,614 |

**Supplementary Table 4. Chloroplast genome assembly statistics of *S. cumini***

| | |
|---|---|
| Genome size | 158,509 bases |
| Size of IR (Inverted Repeat) region | 26,077 bases |
| Size of LSC (Large Single Copy) region | 88,007 bases |
| Size of SSC (Small Single Copy) region | 18,348 bases |
| No. of protein-coding genes | 86 |
| tRNA genes | 37 |
| rRNA genes | 8 |
| GC-content | 37% |

**Supplementary Table 5. Summary statistics of repetitive regions of *S. cumini* genome identified using RepeatMasker**

| | | | | | |
|---|---|---|---|---|---|
| Total length (bases) | | 709,916,230 | | | |
| GC% | | 40.44% | | | |
| Bases masked | | 365,690,999 (51.51%) | | | |
| | | | No. of elements | Length occupied (bp) | % of sequence |
| Retroelements | | | 108,142 | 111,738,717 | 15.74 |
| | SINEs | | 295 | 94,441 | 0.01 |
| | LINEs | | 15,596 | 11,684,105 | 1.65 |
| | | L2/CR1/Rex | 1,009 | 330,328 | 0.05 |
| | | R2/R4/NeSL | 117 | 61,009 | 0.01 |
| | | RTE/Bov-B | 799 | 263,126 | 0.04 |
| | | L1/CIN4 | 13,327 | 10,922,390 | 1.54 |
| | LTR elements | | 92,251 | 99,960,171 | 14.08 |
| | | BEL/Pao | 259 | 84,679 | 0.01 |
| | | Ty1/Copia | 42,835 | 38,095,712 | 5.37 |
| | | Gypsy/DIRS1 | 44,452 | 57,410,705 | 8.09 |
| | | Retroviral | 73 | 33,096 | 0.00 |
| DNA transposons | | | 25,326 | 14,484,075 | 2.04 |
| | hobo-Activator | | 6,684 | 3,892,761 | 0.55 |
| | Tourist/Harbinger | | 1,839 | 1,539,069 | 0.22 |
| Rolling-circles | | | 14,138 | 5,889,064 | 0.83 |
| Unclassified | | | 676,899 | 223,831,938 | 31.53 |
| Total interspersed repeats | | | | 350,054,730 | 49.31 |
| Small RNA | | | 5,138 | 2,342,781 | 0.33 |
| Satellites | | | 168 | 53,950 | 0.01 |
| Simple repeats | | | 135,996 | 6,324,190 | 0.89 |
| Low complexity | | | 20,702 | 1,026,284 | 0.14 |

**Supplementary Table 6. Transcriptome assembly statistics of *S. cumini* species**

| | |
|---|---|
| **Counts of transcripts** | |
| Total trinity 'genes' | 95,459 |
| Total trinity transcripts | 204,525 |
| GC% | 43.91 |
| **Statistics based on all transcript contigs** | |
| Contig N50 | 2,313 |
| Median contig length (bases) | 673 |
| Average contig (bases) | 1,260.52 |
| Total assembled bases | 257,807,839 |
| **Stats based on only longest isoform per 'gene'** | |
| Contig N50 | 1,846 |

| | |
|---|---|
| Median contig length (bases) | 400 |
| Average contig (bases) | 866.66 |
| Total assembled bases | 82,730,644 |

**Supplementary Table 7. KEGG pathways assigned to the coding genes of *S. cumini* species (Pathways with ≥25 genes are mentioned below)**

| KEGG pathways | No. of genes |
|---|---|
| Ribosome | 118 |
| Spliceosome | 97 |
| Prion disease | 87 |
| Protein processing in endoplasmic reticulum | 77 |
| Thermogenesis | 71 |
| Nucleocytoplasmic transport | 70 |
| Cell cycle | 66 |
| Oxidative phosphorylation | 66 |
| Ubiquitin mediated proteolysis | 57 |
| Endocytosis | 55 |
| Ribosome biogenesis in eukaryotes | 55 |
| mRNA surveillance pathway | 51 |
| Nucleotide excision repair | 50 |
| RNA degradation | 49 |
| Purine metabolism | 48 |
| Cysteine and methionine metabolism | 44 |
| MAPK signalling pathway - plant | 40 |
| Glycerophospholipid metabolism | 40 |
| Plant hormone signal transduction | 39 |
| Amino sugar and nucleotide sugar metabolism | 39 |
| Peroxisome | 37 |
| Lysosome | 36 |
| Proteasome | 35 |
| Glycolysis / Gluconeogenesis | 34 |
| Pyruvate metabolism | 34 |
| Homologous recombination | 34 |
| Glycine, serine and threonine metabolism | 33 |
| N-Glycan biosynthesis | 33 |
| Porphyrin metabolism | 33 |
| RNA polymerase | 32 |
| Terpenoid backbone biosynthesis | 31 |
| Starch and sucrose metabolism | 31 |
| DNA replication | 31 |
| Base excision repair | 31 |
| Glycerolipid metabolism | 30 |
| Pyrimidine metabolism | 30 |
| Plant-pathogen interaction | 29 |
| Glyoxylate and dicarboxylate metabolism | 29 |
| Alanine, aspartate and glutamate metabolism | 28 |
| Phagosome | 28 |
| Inositol phosphate metabolism | 27 |

| | |
|---|---|
| Basal transcription factors | 27 |
| Aminoacyl-tRNA biosynthesis | 27 |
| Photosynthesis | 26 |
| mTOR signalling pathway | 26 |
| Various types of N-glycan biosynthesis | 25 |
| Cellular senescence | 25 |

**Supplementary Table 8. COG categories assigned to the coding genes of *S. cumini* species**

| COG categories | No. of genes |
|---|---|
| Function unknown | 13,259 |
| Signal transduction mechanisms | 4,623 |
| Transcription | 3,751 |
| Posttranslational modification, protein turnover, chaperones | 3,726 |
| Carbohydrate transport and metabolism | 2,905 |
| Secondary metabolites biosynthesis, transport and catabolism | 2,421 |
| Replication, recombination and repair | 1,695 |
| Intracellular trafficking, secretion, and vesicular transport | 1,657 |
| Translation, ribosomal structure and biogenesis | 1,645 |
| Amino acid transport and metabolism | 1,491 |
| Energy production and conversion | 1,488 |
| RNA processing and modification | 1,413 |
| Lipid transport and metabolism | 1,394 |
| Inorganic ion transport and metabolism | 1,276 |
| Cell cycle control, cell division, chromosome partitioning | 672 |
| Cell wall/membrane/envelope biogenesis | 595 |
| Cytoskeleton | 560 |
| Coenzyme transport and metabolism | 505 |
| Defense mechanisms | 480 |
| Nucleotide transport and metabolism | 421 |
| Chromatin structure and dynamics | 355 |
| Nuclear structure | 99 |
| Extracellular structures | 73 |
| Cell motility | 5 |

**Supplementary Table 9. No. of *S. cumini* coding genes mapped against publicly available databases**

| Databases | No. of coding genes |
|---|---|
| NCBI-nr | 56,403 (92.17%) |
| Swiss-Prot | 38,046 (62.17%) |
| Pfam-A | 35,733 (58.39%) |
| Overall | 56,532 (92.38%) |

**Supplementary Table 10**. **Inter-species collinearity between *Syzygium* species**

| | No. of collinear blocks | No. of species-specific syntelogs | No. of total collinear genes |
|---|---|---|---|
| ***S. cumini* vs. *S. grande*** | 1,792 | *S. cumini* – 24,890 (40.67%)<br>*S. grande* – 20,425 (51.19%) | 45,315 (44.82%) |
| ***S. aromaticum* vs. *S. grande*** | 469 | *S. aromaticum* – 18,877 (68.57%)<br>*S. grande* – 19,003 (47.62%) | 37,880 (56.18%) |
| ***S. cumini* vs. *S. aromaticum*** | 1,541 | *S. cumini* – 20,919 (34.18%)<br>*S. aromaticum* – 17,594 (63.91%) | 38,513 (43.41%) |

**Supplementary Table 11. KEGG pathways of the *S. cumini* genes present in the inter-species collinear blocks constructed with both *S. aromaticum* and *S. grande* (Pathways with ≥25 genes are mentioned)**

| KEGG pathway | No. of genes |
|---|---|
| Ribosome | 109 |
| Spliceosome | 88 |
| Protein processing in endoplasmic reticulum | 72 |
| Nucleocytoplasmic transport | 66 |
| Cell cycle | 54 |
| Thermogenesis | 54 |
| Endocytosis | 49 |
| Oxidative phosphorylation | 48 |
| Ubiquitin mediated proteolysis | 47 |
| mRNA surveillance pathway | 47 |
| Ribosome biogenesis in eukaryotes | 46 |
| RNA degradation | 44 |
| Cysteine and methionine metabolism | 41 |
| Nucleotide excision repair | 41 |
| Purine metabolism | 40 |
| Plant hormone signal transduction | 39 |
| MAPK signalling pathway - plant | 38 |
| Glycerophospholipid metabolism | 37 |
| Amino sugar and nucleotide sugar metabolism | 36 |
| Glycolysis / Gluconeogenesis | 33 |
| Homologous recombination | 33 |
| Lysosome | 33 |
| Peroxisome | 33 |
| Pyruvate metabolism | 32 |
| Starch and sucrose metabolism | 31 |
| Proteasome | 31 |
| Glycine, serine and threonine metabolism | 30 |
| N-Glycan biosynthesis | 30 |
| Glyoxylate and dicarboxylate metabolism | 28 |
| Glycerolipid metabolism | 27 |
| Pyrimidine metabolism | 27 |
| Porphyrin metabolism | 27 |
| Terpenoid backbone biosynthesis | 27 |
| Base excision repair | 27 |

| | |
|---|---|
| Plant-pathogen interaction | 27 |
| DNA replication | 26 |
| Alanine, aspartate and glutamate metabolism | 25 |
| Basal transcription factors | 25 |
| Aminoacyl-tRNA biosynthesis | 25 |
| Phagosome | 25 |

**Supplementary Table 12. KEGG pathways of the genes included in the species-specific gene clusters of *S. cumini* (Pathways with ≥10 genes are mentioned)**

| KEGG pathway | No. of genes |
|---|---|
| Ribosome | 71 |
| Protein processing in endoplasmic reticulum | 44 |
| Spliceosome | 34 |
| Nucleocytoplasmic transport | 31 |
| Amino sugar and nucleotide sugar metabolism | 27 |
| Plant hormone signal transduction | 27 |
| Glycolysis / Gluconeogenesis | 26 |
| Ubiquitin mediated proteolysis | 26 |
| Starch and sucrose metabolism | 24 |
| mRNA surveillance pathway | 24 |
| Oxidative phosphorylation | 23 |
| Cysteine and methionine metabolism | 23 |
| Peroxisome | 23 |
| Ribosome biogenesis in eukaryotes | 21 |
| RNA degradation | 21 |
| Endocytosis | 21 |
| Cell cycle | 21 |
| Pyruvate metabolism | 20 |
| Lysosome | 20 |
| Glycerophospholipid metabolism | 19 |
| Plant-pathogen interaction | 19 |
| Purine metabolism | 18 |
| Nucleotide excision repair | 17 |
| MAPK signalling pathway - plant | 17 |
| PI3K-Akt signalling pathway | 17 |
| Glyoxylate and dicarboxylate metabolism | 16 |
| Inositol phosphate metabolism | 16 |
| Terpenoid backbone biosynthesis | 16 |
| mTOR signalling pathway | 16 |
| Phagosome | 16 |
| Pentose phosphate pathway | 15 |
| Carbon fixation in photosynthetic organisms | 15 |
| Glycerolipid metabolism | 15 |
| Glycine, serine and threonine metabolism | 15 |
| Porphyrin metabolism | 15 |
| RNA polymerase | 15 |
| AMPK signalling pathway | 15 |
| Aminoacyl-tRNA biosynthesis | 14 |

| | |
|---|---|
| Proteasome | 14 |
| Pyrimidine metabolism | 13 |
| Arginine and proline metabolism | 13 |
| Glutathione metabolism | 13 |
| N-Glycan biosynthesis | 13 |
| Phenylpropanoid biosynthesis | 13 |
| Basal transcription factors | 13 |
| Phosphatidylinositol signalling system | 13 |
| Ascorbate and aldarate metabolism | 12 |
| Various types of N-glycan biosynthesis | 12 |
| Flavonoid biosynthesis | 11 |
| Pentose and glucuronate interconversions | 11 |
| Fructose and mannose metabolism | 11 |
| Galactose metabolism | 11 |
| Methane metabolism | 11 |
| Fatty acid biosynthesis | 11 |
| Fatty acid degradation | 11 |
| Sphingolipid metabolism | 11 |
| Valine, leucine and isoleucine degradation | 11 |
| Tryptophan metabolism | 11 |
| Phenylalanine, tyrosine and tryptophan biosynthesis | 11 |
| Ubiquinone and other terpenoid-quinone biosynthesis | 10 |
| Citrate cycle (TCA cycle) | 10 |
| Alanine, aspartate and glutamate metabolism | 10 |
| Tyrosine metabolism | 10 |
| Phenylalanine metabolism | 10 |
| Pantothenate and CoA biosynthesis | 10 |
| Wnt signalling pathway | 10 |
| Sphingolipid signalling pathway | 10 |
| Cellular senescence | 10 |
| NOD-like receptor signalling pathway | 10 |

**Supplementary Table 13. Highly expanded (>25 expanded genes) annotated gene families**

| | | |
|---|---|---|
| Multidrug resistance protein - MATE family | 'GDSL' lipolytic enzyme family | Feruloyl-CoA 6-hydroxylase (F6H) |
| Methyltransferase | Vacuolar iron transporter homolog | Peroxidase (PER) |
| Shikimate O-hydroxycinnamoyltransferase (HCT) | Protein kinase | Xyloglucan:xyloglucosyl transferase |
| Cinnamoyl-CoA reductase (CCR) | Glutamine synthetase | Cellulose synthase |
| Protein trichome birefringence-like | Flavonol synthase (FLS) | Polygalacturonase |
| Ras-related protein | Monoacylglycerol lipase | MFS transporter, PHS family, inorganic phosphate transporter |
| ABC Transporter | Cytochrome P450 family | Caffeic acid 3-O-methyltransferase (COMT) |

| | | |
|---|---|---|
| Prolyl oligopeptidase family | Cinnamyl-alcohol dehydrogenase (CAD) | AAA ATPase family |
| MADS-box transcription factor | Lipolytic acyl hydrolase (LAH) | Acetylajmaline esterase (AAE) |
| ATP-dependent RNA helicase | E3 ubiquitin-protein ligase | Neomenthol dehydrogenase |

**Supplementary Table 14. KEGG pathways of the *S. cumini* genes included in the highly expanded gene families**

| KEGG pathway | No. of genes |
|---|---|
| Phenylpropanoid biosynthesis | 5 |
| Flavonoid biosynthesis | 5 |
| Endocytosis | 5 |
| Spliceosome | 4 |
| Pentose and glucuronate interconversions | 2 |
| ABC transporters | 2 |
| AMPK signalling pathway | 2 |
| Necroptosis | 2 |

**Supplementary Table 15. KEGG pathways of the positively selected genes in *S. cumini* (Pathways with >5 genes are mentioned)**

| KEGG pathway | No. of genes |
|---|---|
| Ribosome | 17 |
| Glycolysis / Gluconeogenesis | 16 |
| RNA degradation | 16 |
| Ribosome biogenesis in eukaryotes | 15 |
| Cell cycle | 15 |
| Starch and sucrose metabolism | 14 |
| Purine metabolism | 14 |
| Nucleocytoplasmic transport | 14 |
| Protein processing in endoplasmic reticulum | 14 |
| Pyruvate metabolism | 13 |
| Oxidative phosphorylation | 12 |
| Spliceosome | 12 |
| Ubiquitin mediated proteolysis | 11 |
| Proteasome | 11 |
| Endocytosis | 11 |
| Citrate cycle (TCA cycle) | 10 |
| Glyoxylate and dicarboxylate metabolism | 10 |
| Cysteine and methionine metabolism | 10 |
| Aminoacyl-tRNA biosynthesis | 10 |
| Lysosome | 10 |
| Amino sugar and nucleotide sugar metabolism | 9 |
| Glycerophospholipid metabolism | 9 |
| Phagosome | 9 |
| Synaptic vesicle cycle | 9 |
| Valine, leucine and isoleucine degradation | 8 |
| mRNA surveillance pathway | 8 |
| Plant hormone signal transduction | 8 |

| | |
|---|---|
| Peroxisome | 8 |
| Propanoate metabolism | 7 |
| Inositol phosphate metabolism | 7 |
| Carbon fixation in photosynthetic organisms | 7 |
| Pyrimidine metabolism | 7 |
| Ubiquinone and other terpenoid-quinone biosynthesis | 7 |
| Terpenoid backbone biosynthesis | 7 |
| Phenylpropanoid biosynthesis | 7 |
| Drug metabolism - other enzymes | 7 |
| AMPK signalling pathway | 7 |
| Cellular senescence | 7 |
| Pentose and glucuronate interconversions | 6 |
| Sulfur metabolism | 6 |
| Steroid biosynthesis | 6 |
| Glycine, serine and threonine metabolism | 6 |
| Tyrosine metabolism | 6 |
| Basal transcription factors | 6 |
| Nucleotide excision repair | 6 |
| MAPK signalling pathway – plant | 6 |
| Tight junction | 6 |
| Regulation of actin cytoskeleton | 6 |
| Plant-pathogen interaction | 6 |

**Supplementary Table 16. KEGG pathways of the *S. cumini* genes showing unique amino acid substitution with functional impact (Pathways with ≥5 genes are mentioned)**

| KEGG pathway | No. of genes |
|---|---|
| Protein processing in endoplasmic reticulum | 13 |
| Cell cycle | 12 |
| Starch and sucrose metabolism | 11 |
| Plant hormone signal transduction | 11 |
| Glycerophospholipid metabolism | 10 |
| MAPK signalling pathway – plant | 10 |
| Amino sugar and nucleotide sugar metabolism | 9 |
| Peroxisome | 9 |
| Spliceosome | 8 |
| Terpenoid backbone and ubiquinone and other terpenoid-quinone biosynthesis | 7 |
| Glyoxylate and dicarboxylate metabolism | 7 |
| Cysteine and methionine metabolism | 7 |
| Ribosome | 7 |
| Aminoacyl-tRNA biosynthesis | 7 |
| Wnt signalling pathway | 7 |
| Phosphatidylinositol signalling system | 7 |
| Lysosome | 7 |
| Cellular senescence | 7 |
| Plant-pathogen interaction | 7 |
| Glycolysis / Gluconeogenesis | 6 |
| Pyruvate metabolism | 6 |

| | |
|---|---|
| Phenylalanine, tyrosine and tryptophan biosynthesis | 6 |
| Phenylpropanoid biosynthesis | 6 |
| mRNA surveillance pathway | 6 |
| Ubiquitin mediated proteolysis | 6 |
| Proteasome | 6 |
| DNA replication | 6 |
| PI3K-Akt signalling pathway | 6 |
| Endocytosis | 6 |
| Regulation of actin cytoskeleton | 6 |
| NOD-like receptor signalling pathway | 6 |
| Flavonoid biosynthesis | 5 |
| Fructose and mannose metabolism | 5 |
| Inositol phosphate metabolism | 5 |
| Carbon fixation in photosynthetic organisms | 5 |
| Fatty acid degradation | 5 |
| Glycerolipid metabolism | 5 |
| Purine metabolism | 5 |
| Pyrimidine metabolism | 5 |
| Arginine and proline metabolism | 5 |
| Biosynthesis of various plant secondary metabolites | 5 |
| Ribosome biogenesis in eukaryotes | 5 |
| RNA degradation | 5 |
| TGF-beta signalling pathway | 5 |
| cGMP-PKG signalling pathway | 5 |

**Supplementary Table 17. KEGG pathways of the *S. cumini* genes showing higher nucleotide divergence (Pathways with >1 genes are mentioned)**

| KEGG pathway | No. of genes |
|---|---|
| Spliceosome | 7 |
| Ribosome | 3 |
| Oxidative phosphorylation | 3 |
| Phagosome | 3 |
| Synaptic vesicle cycle | 3 |
| Pyrimidine metabolism | 2 |
| Flavonoid biosynthesis | 2 |
| Drug metabolism - other enzymes | 2 |
| Nucleocytoplasmic transport | 2 |
| Protein processing in endoplasmic reticulum | 2 |
| Ubiquitin mediated proteolysis | 2 |
| RNA degradation | 2 |
| Lysosome | 2 |
| Circadian rhythm - plant | 2 |

**Supplementary Table 18. Gene Ontology (GO) categories that were enriched in *S. cumini* MSA genes (TPM > 1) (Top 10 categories are mentioned below)**

| Biological processes | | |
|---|---|---|
| **GO term ID** | **Description** | **p-value** |
| GO:0015748 | Organophosphate ester transport | 0.0073344 |
| GO:0006414 | Translational elongation | 0.014264 |
| GO:0034248 | Regulation of cellular amide metabolic process | 0.020711 |
| GO:0010876 | Lipid localization | 0.027625 |
| GO:0044093 | Positive regulation of molecular function | 0.028080 |
| GO:0016192 | Vesicle-mediated transport | 0.029656 |
| GO:0043900 | Regulation of multi-organism process | 0.030534 |
| GO:0033365 | Protein localization to organelle | 0.036683 |
| GO:0040011 | Locomotion | 0.038810 |
| GO:0006605 | Protein targeting | 0.041498 |
| **Cellular component** | | |
| GO:0030135 | Coated vesicle | 0.013460 |
| GO:0005802 | Trans-Golgi network | 0.017526 |
| GO:0005798 | Golgi-associated vesicle | 0.027701 |
| GO:0005730 | Nucleolus | 0.041986 |
| GO:0005795 | Golgi stack | 0.055409 |
| GO:0030133 | Transport vesicle | 0.064633 |
| GO:0048475 | Coated membrane | 0.080370 |
| GO:0030014 | CCR4-NOT complex | 0.16002 |
| GO:0012506 | Vesicle membrane | 0.16952 |
| GO:1904949 | ATPase complex | 0.17169 |
| **Molecular function** | | |
| GO:0005319 | Lipid transporter activity | 0.0099818 |
| GO:0042578 | Phosphoric ester hydrolase activity | 0.015297 |
| GO:0030276 | Clathrin binding | 0.025551 |
| GO:0016741 | Transferase activity, transferring one-carbon groups | 0.042402 |
| GO:0004386 | Helicase activity | 0.048823 |
| GO:0001871 | Pattern binding | 0.053585 |
| GO:0019899 | Enzyme binding | 0.086583 |
| GO:0008237 | Metallopeptidase activity | 0.10679 |
| GO:0019239 | Deaminase activity | 0.12891 |
| GO:0000049 | tRNA binding | 0.14593 |

**SUPPLEMENTARY FIGURES**

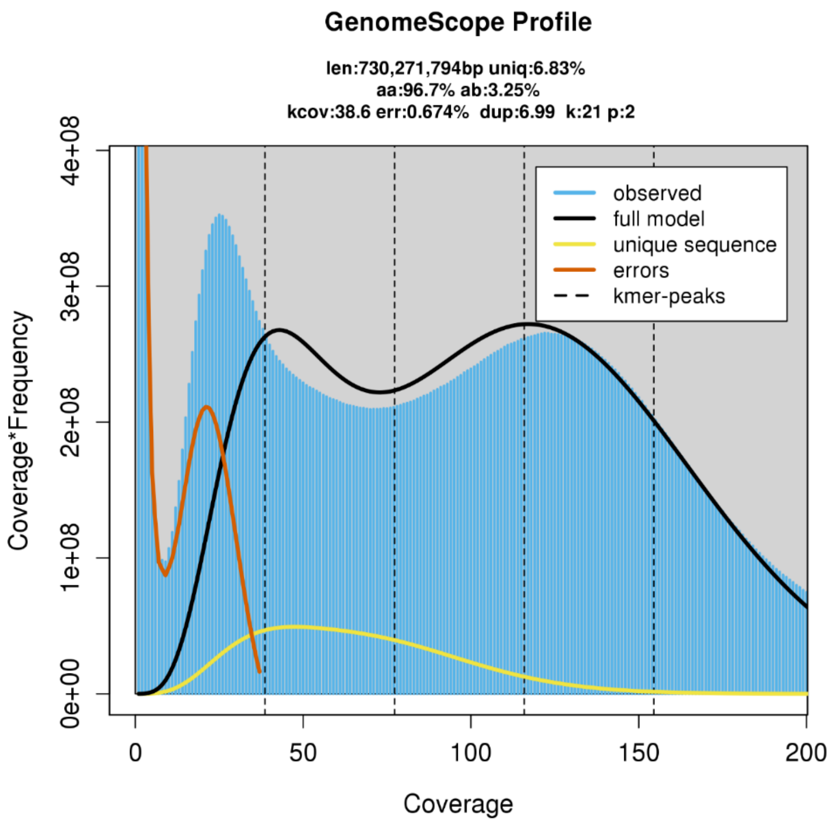


**Supplementary Figure 1.** GenomeScope profile of *S. cumini* genome showing genome size and heterozygosity (using tetraploid model) [1].

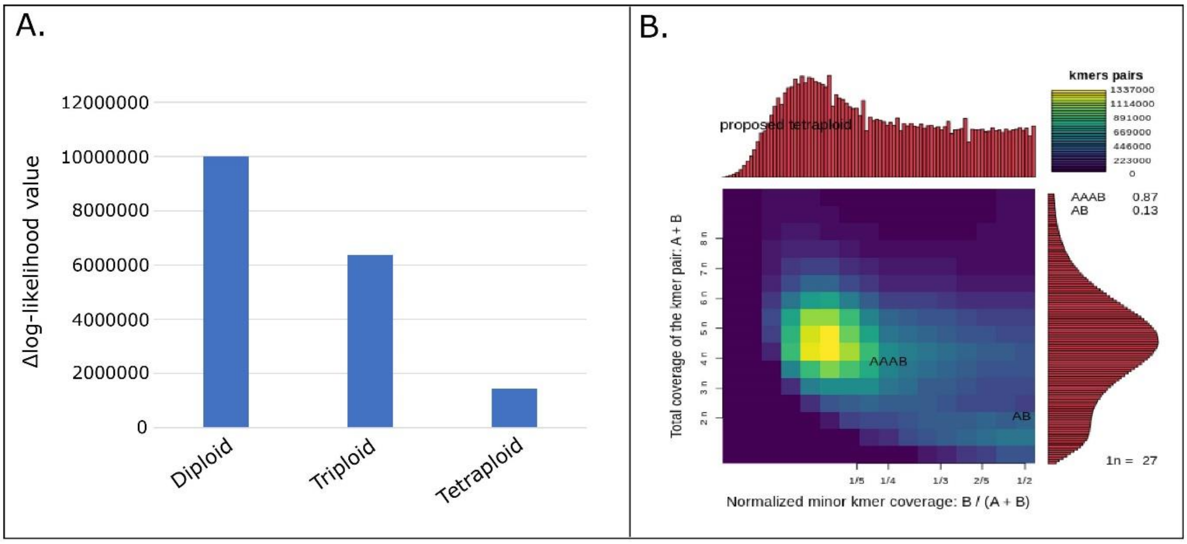


**Supplementary Figure 2.** Ploidy level estimation for *S. cumini* genome. **A.** Δlog-likelihood values for the three fixed models using nQuire [2], **B.** Smudgeplot profile for *S. cumini* genome [1].

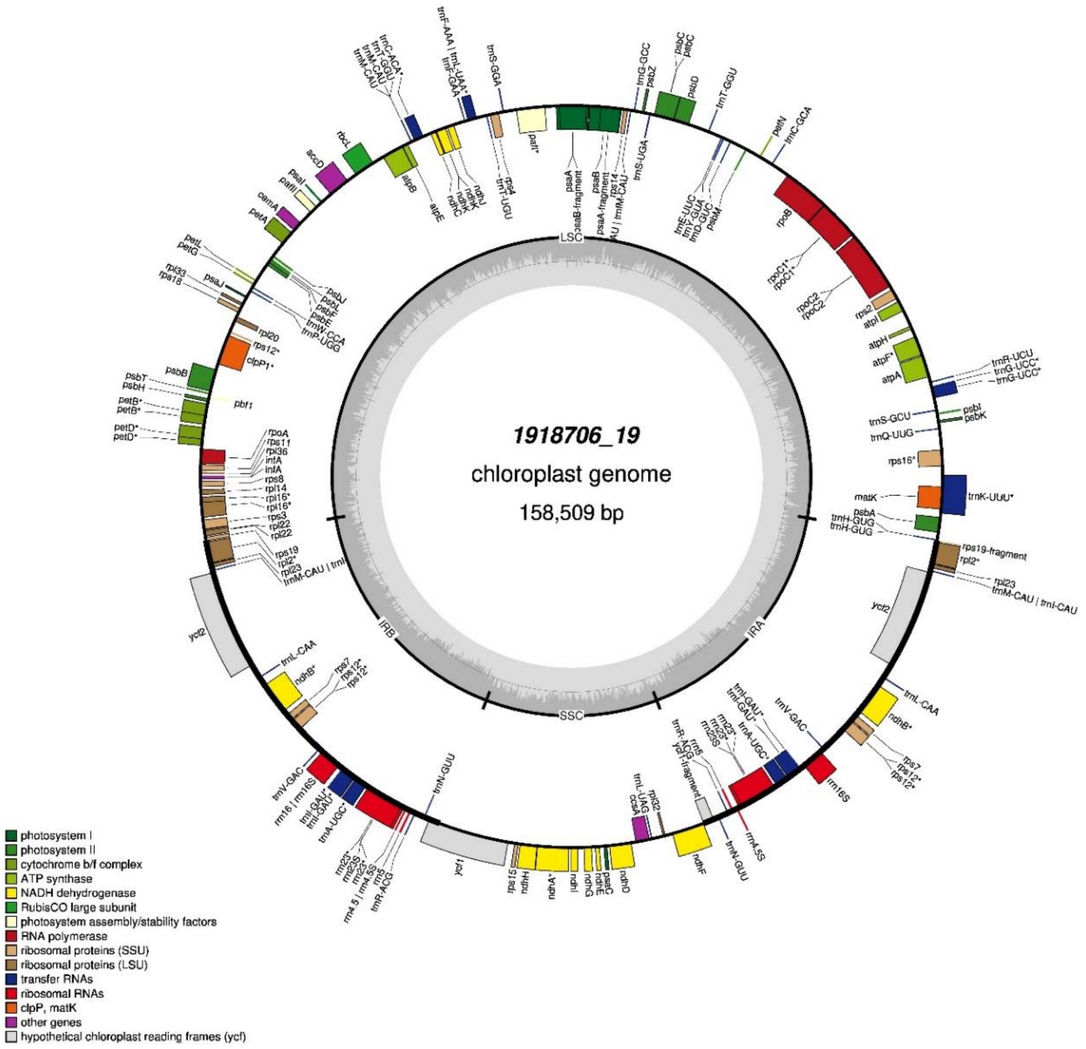


**Supplementary Figure 3**. Chloroplast genome annotation of *S. cumini*

**SUPPLEMENTARY TEXT**

**DNA extraction**

The leaves were collected and cleaned to extract nucleic acid (DNA/RNA). The leaves were taken immediately for RNA extraction. The washed leaves (~3 µg in weight) were homogenized in liquid nitrogen using a pre-cooled autoclaved mortar and pestle. The homogenized leaves were taken in a 50 ml centrifuge tube with 20 mL of Carlson lysis buffer. Carlson lysis buffer was pre-heated at 65˚C for 30 mins. The lysis process was supplemented with the addition of 100 µl of β-mercaptoethanol, 200 µl of Proteinase K (Qiagen, CA, USA), and 100 µl of RNase A (PureLink, ThermoFisher), and incubated at 65˚C for 1 hr with intermittent mixing by inverting the tubes in every 15 mins. After 1 hr, the tube was allowed to cool at room temperature (RT), and subsequently, 100 µl of RNase A was added and incubated at 37˚C for 30 mins for RNA degradation. After RNase treatment, the sample was purified thrice with an equal volume of chloroform: isoamyl alcohol (ratio 24:1) and centrifuged at 4,500xg for 15 mins. The final aqueous phase was collected in a new centrifuge tube and 0.7x ice-cold isopropanol was added. The tube was mixed by inverting it slowly to avoid fragmentation of DNA. The DNA precipitation was facilitated by overnight incubation at room temperature. The room temperature incubation inhibits polysaccharide precipitation along with DNA. The DNA was pelleted down by centrifuging at 5,000xg for 10 mins and washed thrice with 70% ethanol. The DNA pellet was air dried, eluted in 200 µl of nuclease-free water (NFW), and incubated at 37˚C for 10 mins. The quality of DNA was checked on 0.8% agarose gel electrophoresis as well as Nanodrop 8000 spectrophotometer. The DNA was quantified using a qubit ds DNA BR assay kit on Qubit 2.0 fluorometer (Life Technologies, United States).

**Species identification assay**

The extracted DNA was used to amplify two DNA markers: *ITS2* (Internal Transcribed Spacer) and *MatK* (Maturase K). The amplification was performed on Veriti 96 well thermal cycler (Applied Biosystems) using the following primers:

1. *ITS2* forward primer: 5′-GCATCGATGAAGAACGCAGC-3′
   *ITS2* reverse primer: 5′-TCCTCCGCTTATTGATATGC-3′
2. *MatK* forward primer: 5′-CGATCTATTCATTCAATATTTC-3′
   *MatK* reverse primer: 5′-TCTAGCACACGAAAGTCGAAGT-3′

The amplification was evaluated on 2% agarose gel, followed by the purification of amplicons using a PureLink PCR purification kit (Invitrogen, USA). The purified amplicons were sequenced on a Sanger sequencer. The obtained amplicon sequences were aligned against NCBI non-redundant nucleotide database (nt) using BLASTN.

**Genomic sequencing**

The extracted DNA was used to prepare the library on the Chromium controller instrument using Chromium Genome Library and Gel Bead Kit (10x Genomics). The 10x Genomics library was sequenced on Illumina NovaSeq 6000 instrument for generating 150 bp paired-end reads. The DNA was purified using Ampure XP magnetic beads (Beckman Coulter, USA) for Nanopore sequencing. The purified DNA was used to prepare the Nanopore library using SQK-LSK109 and SQK-LSK110 library preparation kit (Oxford Nanopore Technologies, UK). The library was loaded on flowcell and sequenced on a MinION Mk1C sequencer.

**RNA extraction and sequencing**

The RNA was extracted following a similar method that was used for *Syzygium longifolium species* with a few modifications [3]. The RNA precipitation was performed directly using two volumes of 100% ethanol and 1/10 volume of 3M sodium acetate and incubated at 4˚C overnight. The RNA was washed and purified using a RNeasy mini kit (Qiagen, CA, USA). The RNA quality was diluted ten times and was quantified on Qubit 2.0 fluorometer using a qubit ss RNA HS kit (Life Technologies, United States). Quality of the RNA was evaluated using High Sensitivity D1000 ScreenTape on Agilent 2200 TapeStation (Agilent, Santa Clara, CA). The RNA library was prepared using TruSeq Stranded Total RNA Library Preparation kit with the Ribo-Zero Plant workflow (Illumina Inc., CA, USA). The transcriptome library was sequenced to generate 150 bp paired-end reads on Illumina NovaSeq 6000 instrument.

**REFERENCES**


[1] T.R. Ranallo-Benavidez, K.S. Jaron, M.C. Schatz, GenomeScope 2.0 and Smudgeplot for reference-free profiling of polyploid genomes, Nat. Commun. (2020). https://doi.org/10.1038/s41467-020-14998-3.

[2] C.L. Weib, M. Pais, L.M. Cano, S. Kamoun, H.A. Burbano, nQuire: A statistical framework for ploidy estimation using next generation sequencing, BMC Bioinformatics. (2018). https://doi.org/10.1186/s12859-018-2128-z.

[3] J. Soewarto, C. Hamelin, S. Bocs, P. Mournet, H. Vignes, A. Berger, A. Armero, G. Martin, A. Dereeper, G. Sarah, F. Carriconde, L. Maggia, Transcriptome data from three endemic Myrtaceae species from New Caledonia displaying contrasting responses to myrtle rust (Austropuccinia psidii), Data Br. 22 (2019) 794. https://doi.org/10.1016/J.DIB.2018.12.080.
